# Volumetric mechanoplasticity couples melanoma drug tolerance to susceptibility to CD8⁺ T cell killing

**DOI:** 10.64898/2026.01.27.701903

**Authors:** Xingjian Zhang, Dingyao Zhang, Xiangyu Gong, Andrew Peiran Yu, Rakesh K. Jain, Lance L. Munn, Michael Mak

## Abstract

Cell swelling has been reported in physiological tumor contexts, but whether sustained cell volume expansion leaves a durable state that reshapes drug and immune responses in melanoma is unknown. Using controlled hypotonic dilution as an experimental handle, we compare two priming regimens with comparable hypotonic exposure but different magnitude and persistence of volume expansion, enabling us to test which post-recovery phenotypes track with sustained enlargement beyond hypotonic exposure alone. Sustained enlargement produces a measurable size imprint after return to isotonic conditions, accompanied by nuclear lamina and chromatin remodeling with p53 pathway engagement and suppression of replication and growth programs. After recovery, primed cells reinforce F-actin organization, migrate faster in 3D collagen, and induce antioxidant and anti-ferroptosis defenses consistent with improved stress survival. A gene expression signature derived from this response is associated with poorer outcome in TCGA skin cutaneous melanoma. Unexpectedly, the same volume history also increases IFN-γ response, elevates MHC-I, reduces sialylation, and increases susceptibility to CD8^+^ T cell killing. These findings indicate that persistent volume history can couple drug tolerance programs to an exploitable increase in immune visibility.

## Introduction

Advanced melanoma can show marked initial responses to targeted therapy, particularly BRAF and MEK inhibition, yet relapse from minimal residual disease remains common and lethal^1^. Single-cell and spatial profiling further indicate that residual melanoma cells occupy multiple nongenetic states along an inferred differentiation trajectory, including proliferative, invasive, neural crest-like, and undifferentiated programs that can coexist within the same tumor^1–3^. These observations have placed phenotype switching, defined here as transitions among therapy-associated cell programs without new driver mutations, at the centre of resistance, because distinct states differ in sensitivity to MAPK inhibition and to immune-mediated elimination^4^. Consistent with this state dependence, clinical series report that a subset of patients whose tumors progress on BRAF or MEK inhibition can still derive substantial, and sometimes durable, benefit from subsequent immune checkpoint blockade^5,6^ or adoptive T cell therapy^7,8^. Together, these findings motivate a central question: which microenvironmental features bias melanoma toward drug-tolerant states that nevertheless remain susceptible to cytotoxic CD8⁺ T cells?

In parallel with molecular models of plasticity, physical and hydromechanical factors have emerged as a key layer of tumor regulation^9^. Solid tumors accumulate solid stress, elevated interstitial fluid pressure, abnormal stiffness, and disordered architecture, all of which can shape progression and treatment response^9,10^. Measurements in preclinical models and patient samples further connect tumor growth, matrix deposition, cell contractility, and host tissue confinement to increased mechanical stress, while vascular leak and impaired lymphatic drainage contribute to edema and elevated fluid pressure^11–13^. This literature has clarified how these tissue-scale traits compress vessels, reduce perfusion, and hinder drug delivery. It also highlights glycosaminoglycan-rich matrices as major reservoirs for interstitial fluid, suggesting a route by which local fluid handling can reshape transport and mechanics. At the cellular scale, studies in confined microchannels show that directed water permeation and volume regulation can drive migration, indicating that fluid flux and volume control can directly influence cell behavior^14^. Together, these observations motivate a time-integrated view in which tumor hydromechanics may be experienced as a history rather than a snapshot. We therefore asked whether a sustained history of cell volume expansion can durably reprogram melanoma behavior, and we built a controlled priming framework to test this idea.

Recent work in 3D systems has begun to quantify cell volume expansion as an actively regulated feature of dense microenvironments. In mammary cancer organoids, cells at the periphery and invasive front are larger and more dynamic than those in the core, and this swollen subpopulation has been attributed to supracellular fluid flow through gap junctions. Disrupting this intercellular transport suppresses swelling and delays invasion, and analogous gradients in cell and nuclear size have been reported in human breast cancer specimens^15,16^. In a complementary invasion model, coordinated collective volume expansion contributes to forces that stretch and perforate basement membrane, enabling breach without classical invadopodia^17^. Outside cancer, controlled volume changes in soft matrices can couple to fate regulation over multiday timescales, positioning cell volume as a control parameter rather than a passive readout^18,19^. More broadly, multiday mechanical histories can prime durable state shifts across systems, including melanoma^3,20–22^. Together, these studies raise the possibility that defined histories of volume expansion can reshape the set of tumor states that cells occupy and transition between. What remains poorly resolved is how such histories rebalance proliferation versus invasion programs, alter tolerance to cytotoxic therapies, and intersect with the pathways that govern CD8⁺ T cell recognition and killing.

Most current models of melanoma plasticity are framed in terms of transcription factors, epigenetic regulators, metabolic rewiring, and immune signaling pathways, which have been essential for mapping how melanoma escapes targeted therapy^4,23^. Despite this progress, the role of cell volume regulation as a physical determinant of state transitions remains underexplored. Now that emerging 3D evidence indicates that sustained cell volume expansion can be regulated and functionally consequential in dense tissues^15,16^, cell volume becomes a timely variable to interrogate in melanoma plasticity. In general, cell diameter and volume can change through multiple mechanisms, including osmotic and ionic gradients coupled to water transport, regulation of ion channels and transporters, cell-cycle state, cytoskeletal and cortical tension, and physical constraints imposed by adhesion, confinement, and extracellular matrix properties. In tumors, these inputs can vary across space and time and are difficult to isolate in vivo. We therefore use extracellular osmolarity as a controllable experimental handle to impose reproducible hydromechanical histories, not as a direct proxy for any single tumor osmotic state^14,15,17,18,32^. In a tractable B16 melanoma system, we apply two multiday priming regimens that expose cells to comparable hypotonic conditions but produce different magnitude and persistence of volume expansion, allowing us to test which post-recovery phenotypes track with sustained enlargement beyond hypotonic exposure alone. We then quantify early post-recovery cell states after return to isotonic conditions, assess nuclear architecture and chromatin remodeling, measure organelle stress, and test how these histories reshape growth, invasion, and response to cytotoxic therapy. Finally, we derive a volume-linked transcriptional program and evaluate its association with clinical outcome in TCGA skin cutaneous melanoma.

On this basis, we introduce the concept of volumetric mechanoplasticity. We define it as a hydromechanical history in which a defined priming episode induces sustained cell volume expansion and stores a measurable post-recovery state that remains detectable after return to isotonic conditions over the early post-recovery period. Here, volume expansion history refers to prior sustained cell volume expansion during priming together with its post-recovery imprint, not the instantaneous volume change itself. In this model, volume expansion history biases melanoma toward a slow-cycling, invasion-competent program coupled to stress adaptation that increases tolerance to cytotoxic therapy. Unexpectedly, the same history also reshapes immune visibility phenotypes. Together, these findings position volume expansion history as a cell-intrinsic hydromechanical variable that can align drug tolerance with exploitable CD8^+^ T cell susceptibility within specific melanoma cell states, offering a physical explanation for how resistance and immune sensitivity can coexist within the same tumor ecosystem.

## Results

### Volume expansion history is associated with a durable post-recovery size imprint

Melanoma can exhibit substantial size heterogeneity, but in patient tissue the key question is whether size variation reflects organized, population-level structure rather than isolated single-cell differences. We quantified nuclear area in a representative human melanoma section as an image-based proxy for cellular volume^15,18,21,24^, segmenting nuclei with Cellpose-SAM^25,26^, a SAM-based instance segmentation method, and mapping log₂-normalized nuclear area across the tissue (Fig. 1A). Because nuclear cross-sections in fixed tissue are influenced by sectioning geometry^27^, heterogeneous cell-cycle state^28^, and rare intrinsically enlarged nuclei^29^, we did not interpret the global size distribution. Global Moran’s I^30^ indicated significant positive spatial autocorrelation of nuclear area across the section, while local Moran’s I (LISA)^31^ identified High–High and Low–Low clustered domains together with High–Low and Low–High spatial outliers (Extended Data Fig. 1). In parallel, we annotated representative neighborhoods on the heatmap and quantified their normalized nuclear area distributions, revealing regions enriched for relatively small, intermediate, or expanded nuclei (Fig. 1B). These observations establish that melanoma can organize into local nuclear-size niches, motivating a controllable system to test whether defined volume expansion histories encode durable post-recovery phenotypes.

**Figure 1.**
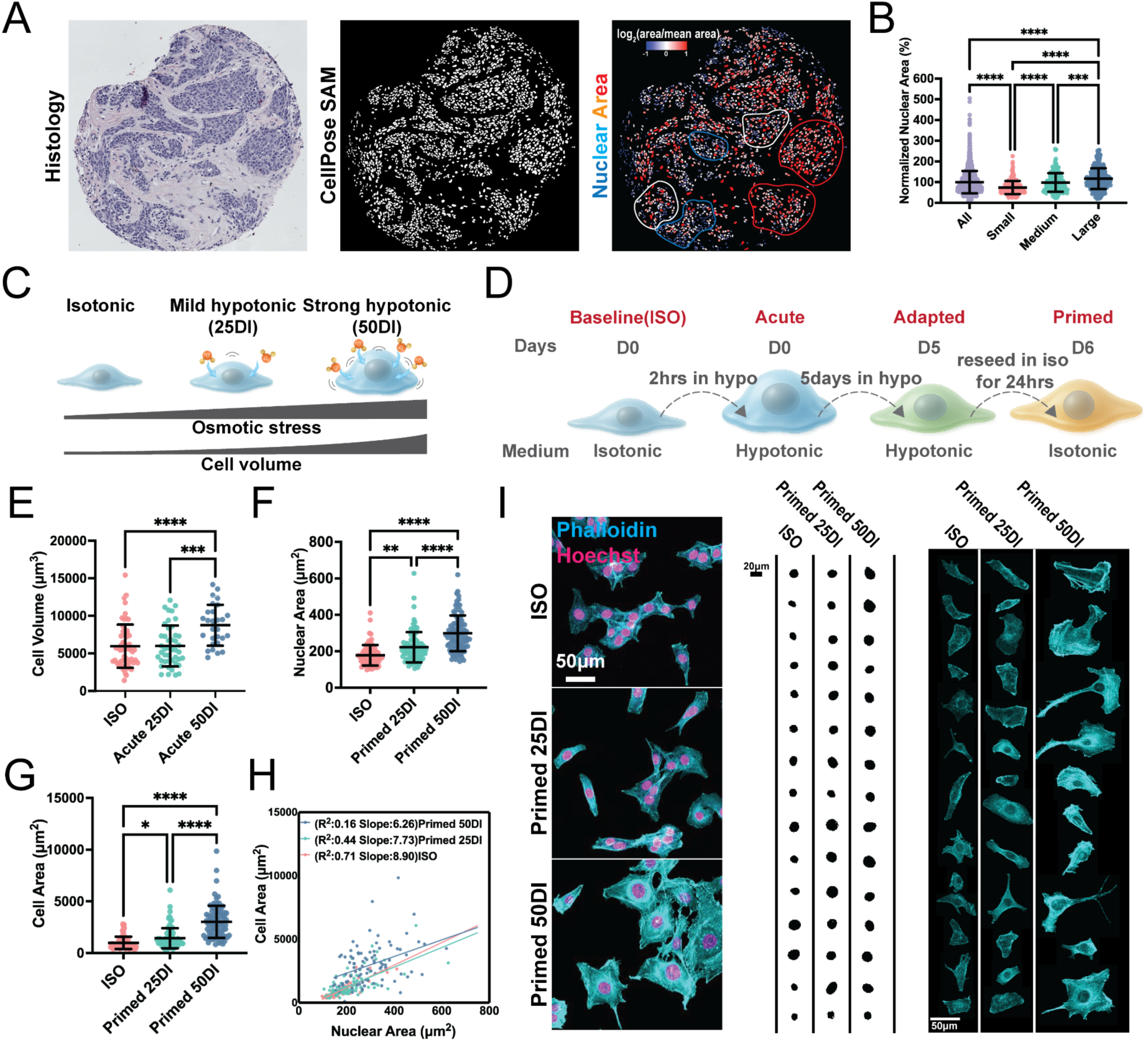
Volume expansion history is associated with a durable post-recovery size imprint. (A) Human melanoma section showing histology, nuclear segmentation, and a nuclear area heatmap displaying log₂(nuclear area divided by mean nuclear area). Outlines mark representative spatial neighborhoods enriched for relatively small, intermediate, or large nuclei. Spatial autocorrelation and data driven segmentation analyses supporting nonrandom neighborhood structure are provided in Extended Data Fig. 1. (B) Quantification of normalized nuclear area across all nuclei and within the annotated neighborhoods. Each dot represents one nucleus (n = 2261 total nuclei, n ≥ 188 per neighborhood). (C) Conceptual schematic defining two hydromechanical histories using extracellular dilution: an osmotic history (25DI) that imposes hypotonic stress with minimal acute swelling in the primary model, and a volumetric history (50DI) that induces robust acute swelling and sustained enlargement during multiday adaptation. (D) Experimental timeline defining isotonic baseline (ISO), acute exposure (2 h), adaptation (5 days in hypotonic medium), and the primed state defined as reseeding into ISO followed by 24 h recovery. (E) Acute single cell volume response in B16F0 across ISO, acute 25DI, and acute 50DI after 2 h (n ≥ 28 cells per condition, 2 independent experiments). Cross line acute swelling and shape responses and osmolality calibration are shown in Extended Data Fig. 2. (F) Single cell quantification of projected nuclear area in primed cells across ISO, primed 25DI, and primed 50DI (n ≥ 76 cells per condition, 3 independent experiments). (G) Single cell quantification of projected cell area in primed cells across ISO, primed 25DI, and primed 50DI (n ≥ 76 cells per condition, 3 independent experiments). (H) Single cell coupling between nuclear area and cell area across ISO, primed 25DI, and primed 50DI. Each point represents one cell, and lines indicate linear fits for visualization of scaling and condition shifts. (I) Representative confocal images of primed cells stained for F actin (phalloidin) and nuclei (Hoechst) across ISO, primed 25DI, and primed 50DI, with segmented nuclear masks and cropped single cell F actin images. Scale bars as indicated. Scatter plots show mean ± s.d.

To create a controllable multiday volume trajectory, we used extracellular dilution of culture medium with deionized water as an experimental handle to modulate hypotonic load and the associated swelling response^14,15,18,32^. We operationally contrasted two priming histories that share sustained hypotonic exposure but differ in the magnitude of the early swelling response. In the primary system, 25DI imposed hypotonic stress with weaker acute swelling, whereas 50DI induced robust acute swelling that persisted during multiday adaptation (Fig. 1C; Extended Data Fig. 2A). We staged this trajectory into acute exposure (2 h), multiday adaptation (5 days), and a primed readout defined as 24 h after reseeding into isotonic medium (Fig. 1D). Consistent with this design, acute measurements showed a strong volume increase under 50DI, whereas 25DI produced a smaller acute volume gain that was not readily detectable under our sampling in B16F0 cells (Fig. 1E). Across melanoma backgrounds, 50DI consistently induced swelling, but YUMM and YUMMER1.7 also shifted toward higher sphericity (Extended Data Fig. 2C–F), which reduced stable spreading and adhesion area and limited multiday priming for downstream imaging-based phenotyping. By contrast, B16F0 maintained a spread morphology under these conditions, supporting its use as the primary model for mechanoplastic assays.

After return to isotonic conditions, the volumetric history produced a pronounced post-recovery size imprint. Primed 50DI cells remained enlarged at both the nuclear and cellular level relative to ISO, whereas primed 25DI cells showed only modest shifts (Fig. 1F,G,I; Extended Data Fig. 3). We next asked whether the nuclear enlargement reflected proportional enlargement of the whole cell rather than a nucleus-specific effect. At the single-cell level, nuclear area and cell area co-varied across conditions, and primed 50DI cells occupied a shifted nuclear–cell size relationship consistent with coordinated enlargement (Fig. 1H). We use this persistent post-recovery enlargement, measured 24 h after reseeding into isotonic medium, as an operational definition of volumetric mechanoplasticity in the early post-recovery window.

### Volumetric history is encoded by a 50DI-enriched adapted transcriptional program with deeper proliferative suppression and broader stress signaling

Building on the osmotic (25DI) and volumetric (50DI) histories defined above, we asked how these distinct perturbations are reflected in the adapted transcriptome. We performed RNA sequencing on cells after 5 days in hypotonic medium and compared each condition to ISO. Adapted 50DI exhibited a markedly larger transcriptional shift than adapted 25DI (Fig. 2A,B), driven by both a greater number of significantly changed genes and a wider set of changes that were specific to 50DI. Consistent with this, gene-level shift scatter indicated that many genes altered in 50DI were not detectably changed in 25DI, rather than following a simple scaled version of the 25DI response (Fig. 2C). Together, these analyses support the interpretation that the volumetric history engages a distinct adapted transcriptional program.

**Figure 2.**
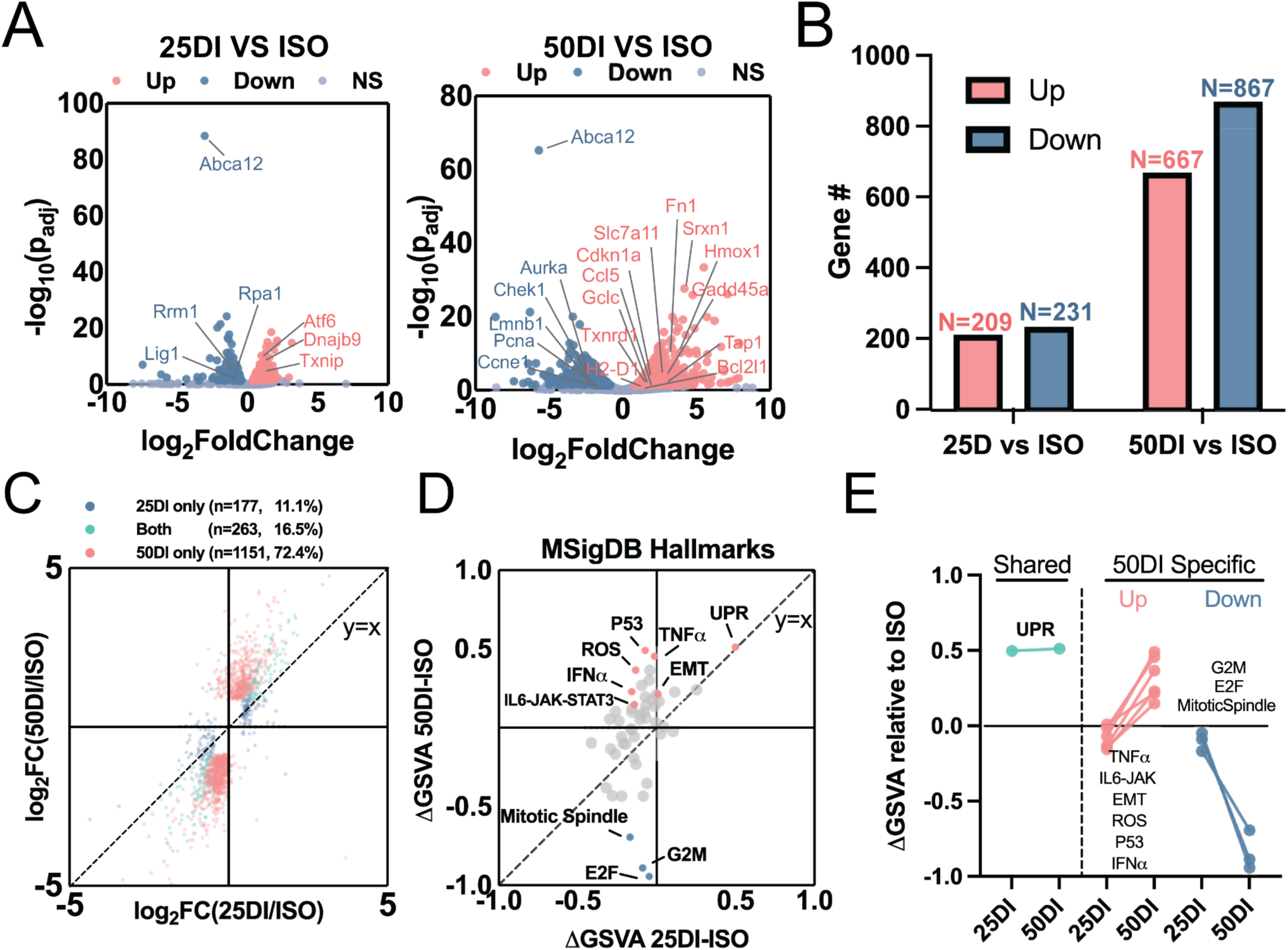
Volumetric history is encoded by a 50DI-enriched adapted transcriptional program with deeper proliferative suppression and broader stress signaling. (A) Volcano plots of differential gene expression comparing adapted 25DI versus ISO (left) and adapted 50DI versus ISO (right). Points represent genes colored by direction and significance relative to ISO using DESeq2 adjusted P value thresholds. Selected representative genes are labeled. (B) Counts of significantly upregulated and downregulated genes in adapted 25DI versus ISO and adapted 50DI versus ISO, highlighting the larger transcriptional shift induced by 50DI. (C) Gene shift scatter comparing log_2_ fold change in adapted 25DI versus ISO (x axis) and adapted 50DI versus ISO (y axis) for genes significant in at least one comparison. Genes are categorized as 25DI only, 50DI only, or significant in both comparisons based on adjusted P value thresholds. (D) Hallmark pathway shift scatter showing ΔGSVA scores for MSigDB Hallmark gene sets. Each point represents a gene set and axes represent ΔGSVA (25DI minus ISO) and ΔGSVA (50DI minus ISO). Selected modules are annotated to highlight a shared stress core and 50DI enriched programs prioritized for downstream functional tests. (E) Summary slope plot of selected modules from panel D, grouped into shared programs and 50DI enriched programs, showing ΔGSVA relative to ISO for adapted 25DI and adapted 50DI. All RNA sequencing analyses were performed on adapted cells (5-day exposure) and compared to ISO.

To summarize shared versus 50DI-enriched modules, we scored pathway activity using Gene Set Variation Analysis (GSVA)^33^ with the MSigDB Hallmark collection^34^ (Fig. 2D,E). Both 25DI and 50DI shared a stress core dominated by unfolded protein response (UPR) programs. In contrast, adapted 50DI showed deeper suppression of proliferative control modules, including E2F targets, G2M checkpoint, and mitotic spindle, together with stronger activation of stress and signaling modules including Reactive Oxygen Species (ROS) pathway, TNFα-NFκB, IL6-JAK-STAT3, epithelial-mesenchymal transition (EMT), p53 pathway, and IFNα response (Fig. 2D,E). We highlight these modules because they provide concrete anchors for downstream phenotypic tests, while a broader reference is provided in Extended Data Fig. 4.

Gene Ontology enrichment provided an orthogonal view consistent with the GSVA structure. Upregulated terms converged on endoplasmic reticulum stress (ER) and protein quality control processes shared across conditions, whereas downregulated terms revealed a large cluster of proliferative control processes that was selectively suppressed in adapted 50DI (Extended Data Fig. 5). In line with this pattern, replication-associated gene sets showed strong negative enrichment in adapted 50DI, and core replication factors were broadly reduced in heatmap view (Extended Data Fig. 6) using Gene Set Enrichment Analysis (GSEA)^35^. Together, these analyses place adapted 25DI as an osmotic stress-biased program and adapted 50DI as a volumetric mechanoadaptive program characterized by stronger proliferative suppression and broader stress-linked pathway engagement.

### Volumetric history is associated with acute mitochondrial stress and persistent nuclear remodeling with p53 engagement

The adapted transcriptomes indicated that 50DI engages stress programs beyond the ER/UPR modules shared with 25DI, prompting us to test whether these signatures align with organelle stress and durable nuclear remodeling. A representative stress gene panel showed broader induction of mitochondrial and oxidative stress associated transcripts in adapted 50DI, alongside ER stress and UPR programs shared across adapted 25DI and 50DI (Fig. 3A). Pathway enrichment recapitulated this structure: ER stress and UPR gene sets were enriched in both adapted states, whereas mitochondrial depolarization and ROS pathways showed a stronger signal in adapted 50DI (Extended Data Fig. 7A–D).

**Figure 3.**
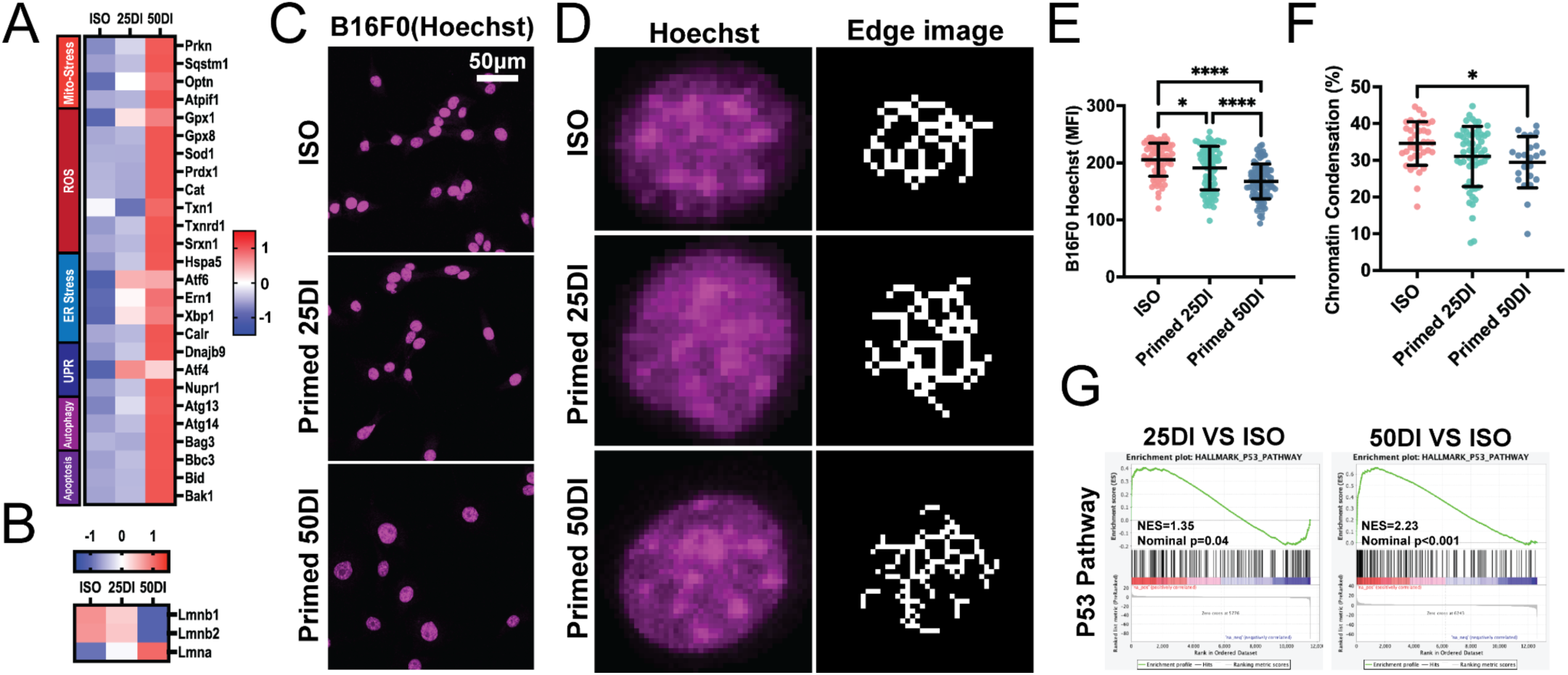
Volumetric history is associated with acute mitochondrial stress and persistent nuclear remodeling with p53 engagement. (A) Heatmap of representative genes associated with mitochondrial stress, ROS, ER stress, unfolded protein response, autophagy, and apoptosis across ISO, adapted 25DI, and adapted 50DI from RNA sequencing. Values are displayed as z scored expression across samples. (B) Heatmap of nuclear lamina genes across ISO, adapted 25DI, and adapted 50DI from RNA sequencing, highlighting a shift toward lower Lmnb1 and higher Lmna in adapted 50DI. Values are displayed as z scored expression across samples. (C) Representative confocal images of primed B16F0 nuclei stained with Hoechst across ISO, primed 25DI, and primed 50DI. Scale bar as indicated. (D) Example single nucleus Hoechst images and the corresponding edge based binary masks used to compute a chromatin condensation index in primed B16F0 cells. (E) Nucleus quantification of mean Hoechst intensity in primed B16F0 nuclei across ISO, primed 25DI, and primed 50DI (n ≥ 76 from 3 independent experiments). (F) Chromatin condensation index in primed B16F0 nuclei across ISO, primed 25DI, and primed 50DI (n ≥ 23 nuclei per condition from 3 independent experiments). (I) GSEA for the Hallmark p53 pathway comparing adapted 25DI versus ISO and adapted 50DI versus ISO. Scatter plots show mean ± s.d. All RNA sequencing analyses were performed on adapted cells (5 day exposure) and compared to ISO.

This stress configuration coincided with nuclear remodeling features that persisted after return to isotonic medium. In the adapted state, lamina transcripts shifted toward lower Lmnb1 and higher Lmna, with the strongest shift in adapted 50DI (Fig. 3B). After recovery, primed 50DI nuclei displayed more diffuse Hoechst patterns relative to ISO (Fig. 3C), motivating a texture-based quantification of chromatin packing using an edge-based condensation metric^37,38^ (Fig. 3D). Across nuclei acquired, primed 50DI showed reduced mean Hoechst intensity and reduced chromatin condensation relative to ISO, whereas primed 25DI showed smaller shifts (Fig. 3E,F). A second melanoma line supported the same directionality, as primed B16F10 nuclei showed reduced DAPI sum intensity after 50DI priming (Extended Data Fig. 8A,B). Consistent with this structural phenotype, a chromosome condensation gene set was more negatively enriched in adapted 50DI (Extended Data Fig. 8C). Because stress and chromatin perturbations of this type can engage p53 signaling^39,40^, we asked whether p53 pathway activity differed across histories. p53 pathway enrichment was stronger in adapted 50DI than in adapted 25DI (Fig. 3G), and stress and p53 target genes showed coherent induction in adapted 50DI (Extended Data Fig. 8D). Together, these results define a volumetric-history associated state in which stronger mitochondrial and oxidative stress signatures align with a lamina shift and a durable post-recovery chromatin decondensation footprint, accompanied by increased p53 pathway engagement.

### Volumetric mechanoplasticity is associated with a sustained reduction in net population expansion that persists after return to isotonic conditions

To translate the transcriptomic suppression of proliferative control into a measurable phenotype, we quantified population expansion during the 5-day hypotonic adaptation period and after reseeding into isotonic medium. During adaptation, brightfield imaging showed progressive divergence in culture density in 50DI relative to ISO, whereas 25DI remained closer to ISO across the time course (Fig. 4A). Direct cell counts confirmed this separation. Adapted 50DI accumulated cells markedly more slowly than ISO and adapted 25DI, both across the full trajectory and in the day 5 over day 0 fold-change summary (Fig. 4B,C). Consistent with these population-level differences, cell-cycle regulators spanning G1–S, G2–M, and mitotic programs were broadly reduced in the adapted state, with the strongest repression in 50DI (Fig. 4D), and Hallmark GSEA independently showed deeper negative enrichment of E2F targets, G2M checkpoint, and mitotic spindle modules in adapted 50DI (Extended Data Fig. 9). We next asked whether this growth restraint persists after return to isotonic conditions. Upon reseeding into isotonic medium in a 3D collagen assay, primed 50DI cultures expanded more slowly than ISO controls over the first 36 h, as quantified by time-resolved area coverage and by the total cancer-cell occupied area at 36 h (Fig. 4E–G). Because these readouts report net expansion, we use them here to operationally define a durable reduction in proliferative capacity that emerges during volumetric adaptation and remains detectable in the early post-recovery window.

**Figure 4.**
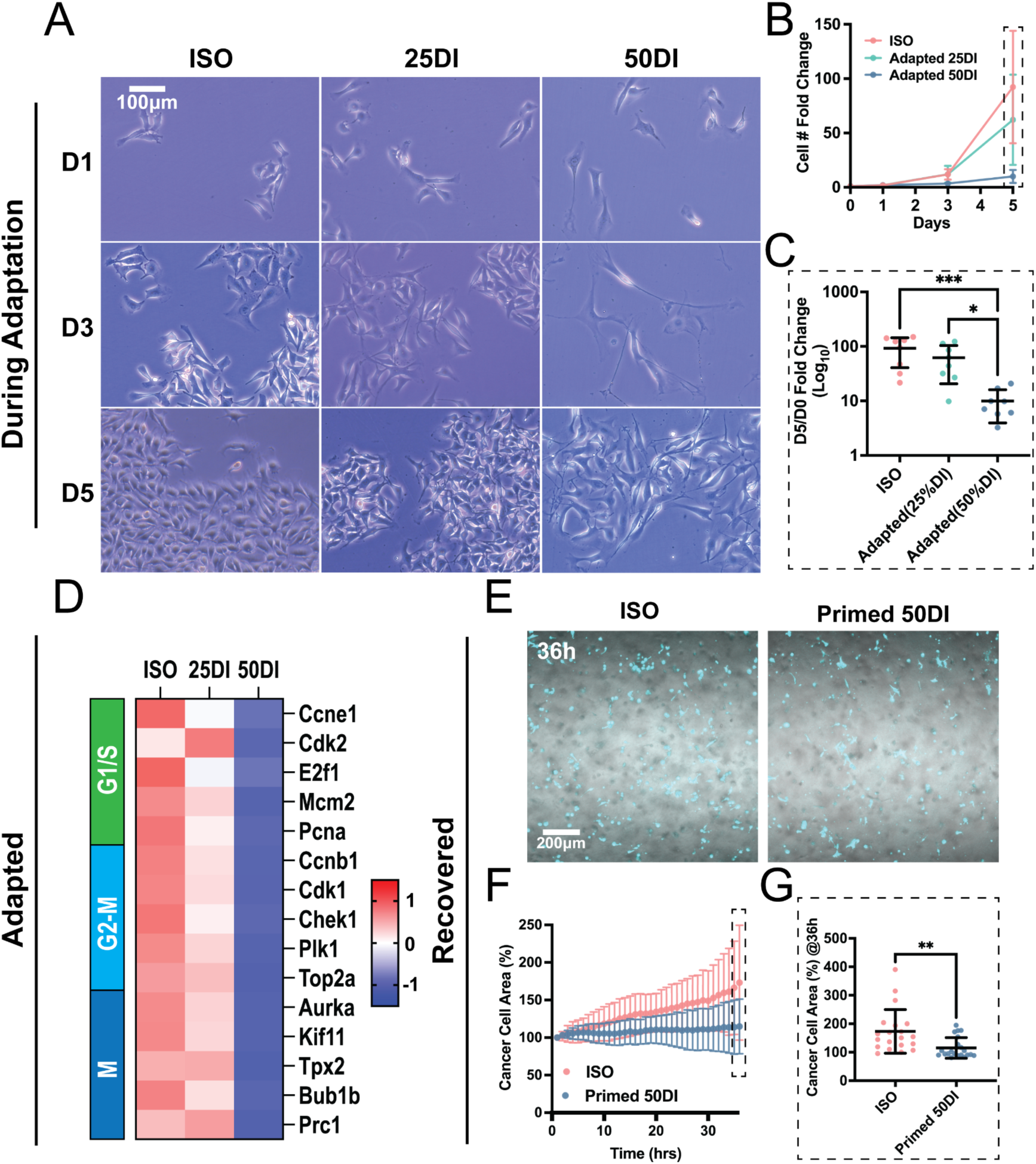
Volumetric mechanoplasticity is associated with a sustained reduction in net population expansion that persists after return to isotonic conditions. (A) Brightfield images showing B16F10 morphology during adaptation in ISO, 25DI, and 50DI at day 1, day 3, and day 5. (B) Cell number fold change over five days in ISO, adapted 25DI, and adapted 50DI. (C) Day 5 over day 0 fold change (log10) in each condition (n = 8). (D) Heatmap of representative G1S, G2M, and M phase regulators across ISO, adapted 25DI, and adapted 50DI from RNA sequencing. Heatmap color bars show z scored expression. (E) Post recovery confocal images of ISO and primed 50DI B16F10 monolayers during isotonic regrowth, with cancer cell masks shown in cyan. (F) Time course of percent area covered by cancer cells over 36 h in ISO and primed 50DI. (G) Cancer cell area at 36 h in ISO and primed 50DI (n ≥ 20 images per condition from 4 independent experiments). Scatter plots show mean ± s.d. All RNA sequencing analyses were performed on adapted cells (5 day exposure) and compared to ISO.

### Volumetric mechanoplasticity shifts melanoma toward an invasion-associated post-recovery state

The proliferative restraint established by volumetric priming raised a critical question: after recovery, are primed cells simply slowed, or do they adopt a distinct functional program. Melanoma can toggle between growth-biased and migration-biased programs^1,41,42^. We therefore asked whether hydromechanical history reweights adhesion and cytoskeletal regulators in a direction consistent with invasion-associated behavior. In the adapted transcriptome, EMT- and adhesion-related genes showed directional remodeling that was most pronounced in 50DI (Fig. 5A). This pattern did not reflect a uniform increase, but rather coordinated gains and losses across modules. Fn1 and Col1a1 were induced, consistent with activation of an interstitial extracellular matrix gene program enriched for fibronectin and type I collagen components^43–45^. In parallel, integrin expression shifted away from laminin-associated α6β4 and toward fibronectin-associated α5β1: Itga6 and Itgb4 were reduced, whereas Itga5 and Itgb1 increased, with the clearest shifts in 50DI (Fig. 5A), consistent with a transition away from laminin-anchored adhesion programs^46^ and toward fibronectin-linked engagement^47,48^. Contractility-linked signaling was also selectively elevated in 50DI, with increased Rhoa, Rhob, and Rhoc together with actomyosin components Acta2 and Myh9^12,49–51^. The epithelial junction marker Cdh1 was reduced, consistent with weakened cell–cell adhesion^52^. Hallmark analysis independently supported EMT enrichment in both adapted 25DI and adapted 50DI relative to ISO (Fig. 5B), with the broader adhesion and contractility remodeling most evident under 50DI.

**Figure 5.**
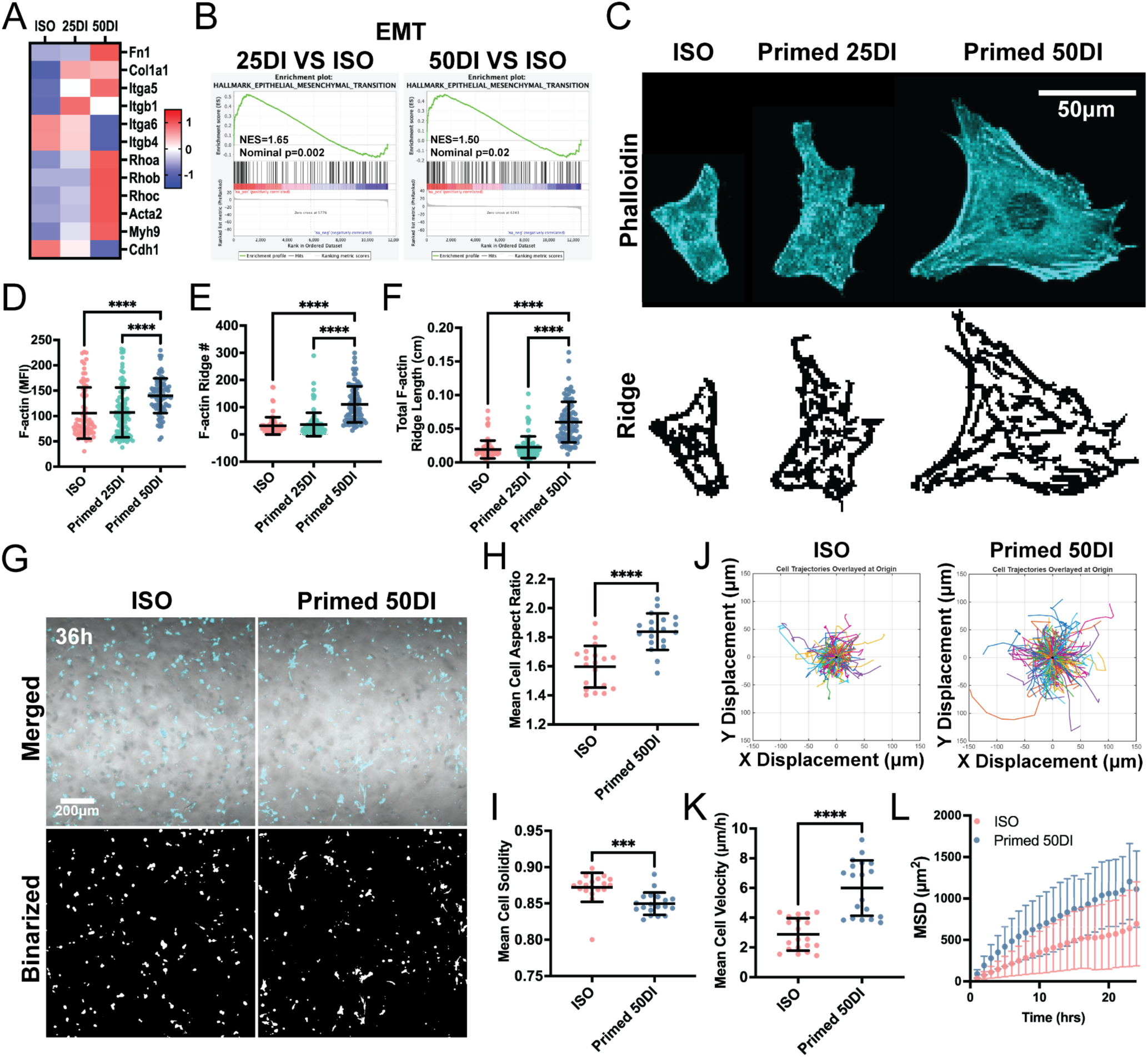
Volumetric mechanoplasticity shifts melanoma toward an invasion-associated post-recovery state. (A) Heatmap of representative EMT and adhesion related genes across ISO, adapted 25DI, and adapted 50DI from RNA sequencing. Heatmap color bars show z scored expression. (B) GSEA for the Hallmark epithelial mesenchymal transition gene set in adapted 25DI (left) and adapted 50DI (right) relative to ISO. (C) Post recovery phalloidin staining of ISO, primed 25DI, and primed 50DI cells, with corresponding ridge based F-actin skeletons shown below. (D–F) Single cell quantification of F-actin organization including stress fiber mean fluorescence intensity (D), ridge number per cell (E), and total ridge length per cell (F) (n≥76 cells per condition from 3 independent experiments). (G) Representative fields at 36 h in 3D collagen showing merged overlays with cyan cancer cell masks (upper) and binarized segmentation masks (lower) for ISO and primed 50DI cultures. (H–I) Single cell morphology metrics at 36 h including mean cell aspect ratio (H) and solidity (I) for ISO and primed 50DI cultures (n≥18 from 4 independent experiments). (J) Representative x–y trajectories of individual cells overlaid at the origin for ISO and primed 50DI cultures. (K) Single cell mean migration velocity for ISO and primed 50DI cultures (n≥18 per condition from 4 independent experiments).(L) Mean squared displacement as a function of time for ISO and primed 50DI cultures. Scatter plots and curves show mean ± s.d. All RNA sequencing analyses were performed on adapted cells (5 day exposure) and compared to ISO.

These transcriptional shifts motivated a structural test after return to isotonic conditions. Following recovery, phalloidin staining revealed a reinforced stress fiber network in primed 50DI cells (Fig. 5C). Ridge-based skeletonization captured this remodeling at single-cell resolution by extracting a stress fiber ridge network from the F-actin signal (Fig. 5C). Quantification showed that primed 50DI increased F-actin intensity, ridge number, and total ridge length relative to ISO (Fig. 5D–F). This reinforcement was specific to 50DI, as primed 25DI remained closer to ISO. Reinforced stress fibers and expanded actin ridge networks are commonly associated with stronger traction and more persistent migration in two-dimensional and three-dimensional settings^53,54^. We therefore interpret this post-recovery actin architecture as a structural marker consistent with an invasion-associated state.

Because the dominant transcriptional remodeling and the post-recovery cytoskeletal reinforcement were most evident under 50DI, we focused motility analyses on this volumetric mechanoplastic state. Actomyosin remodeling is closely linked to migration in three-dimensional environments^54^, motivating a test of post-recovery behavior in 3D collagen. We therefore assessed single-cell motility in 3D collagen after recovery. At 36 h, primed 50DI cultures displayed cell shapes consistent with an invasion-associated motility phenotype. Using two-dimensional shape metrics extracted from projected x–y cell outlines in the 3D collagen assay, aspect ratio increased and solidity decreased relative to ISO controls (Fig. 5G–I). Here, solidity is defined as the ratio of the projected cell area to the area of its convex hull (Area/Convex Area), such that lower values indicate a less convex, more irregular or protrusive outline. Trajectory overlays showed broader dispersal from the origin (Fig. 5J), and mean velocity together with mean squared displacement (MSD) supported increased displacement dynamics in primed 50DI cultures (Fig. 5K,L). This effect was preserved in hyaluronan-containing collagen matrices (Extended Data Fig. 10), a context that better approximates glycosaminoglycan-rich tumor stroma found in many solid tumors^9,12^.

### Volumetric mechanoplasticity links an invasion-associated phenotype to stress-tolerant survival programs

The invasion-associated post-recovery phenotype raised a practical question: does volumetric history also engage stress-tolerance programs that could increase resistance to cytotoxic therapy. To begin addressing this, we asked whether the adapted 50DI transcriptome activates coherent survival modules. Using resistance module scoring, we observed coordinated activation selectively in adapted 50DI, whereas adapted 25DI remained closer to ISO (Fig. 6A). The strongest gains aligned with antioxidant and glutathione handling^55,56^, anti-ferroptosis defense^55,57^, and IL6-JAK-STAT3 signaling^58^. Apoptosis regulation also shifted in a direction consistent with reduced commitment to caspase-dependent death^59^. These module-level patterns were supported by representative pathway enrichment (Extended Data Fig. 11A). Together, these data indicate that volumetric history engages multiple layers of stress-response and survival programs rather than a single dominant pathway.

**Figure 6.**
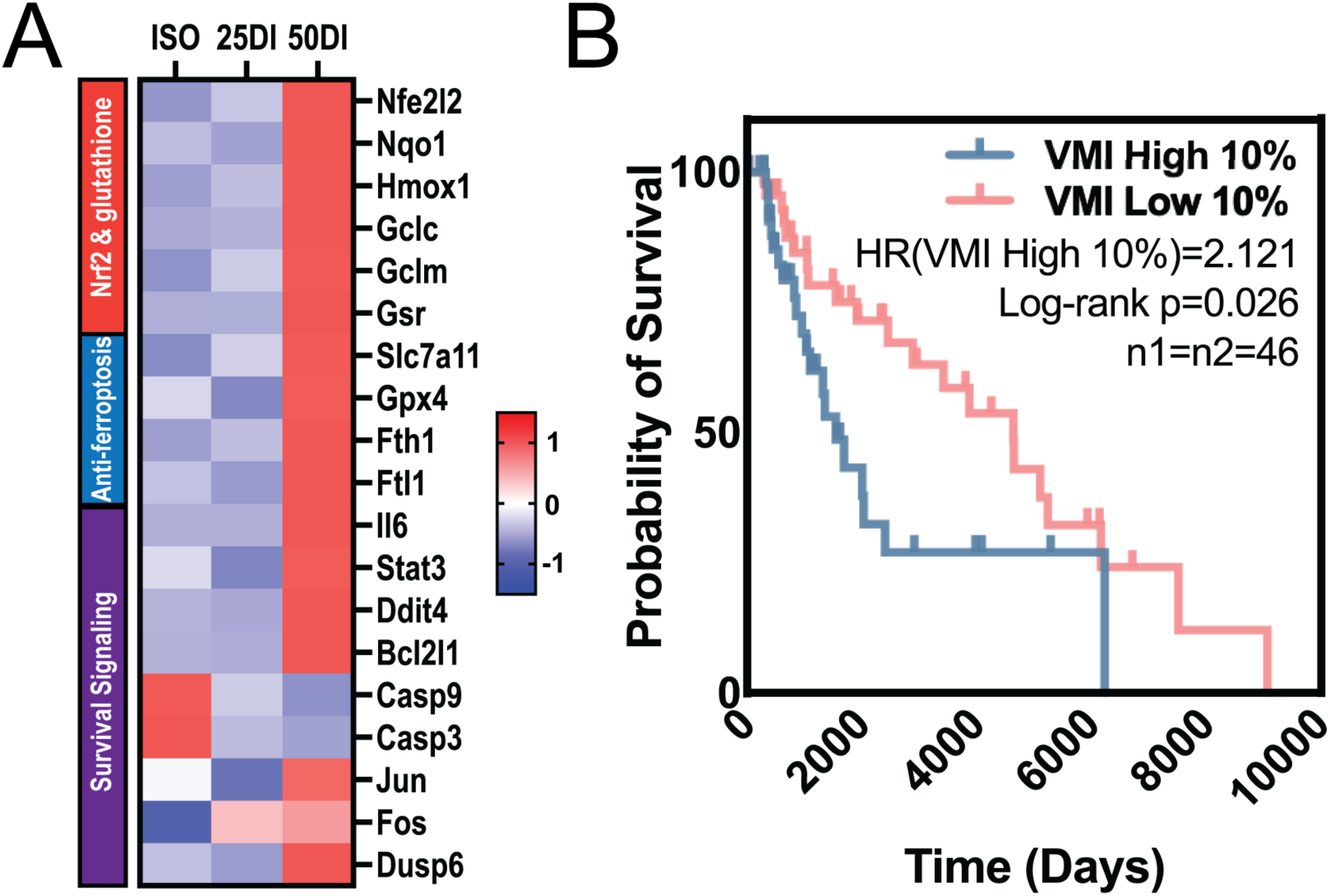
Volumetric mechanoplasticity links an invasion-associated phenotype to stress-tolerant survival programs. (A) Heatmap of curated resistance module activity across ISO, adapted 25DI, and adapted 50DI from RNA sequencing, including Nrf2 dependent glutathione metabolism, anti-ferroptosis defense, IL6-JAK-STAT3 signaling, and apoptosis regulation. Module activity was computed by gene set scoring on normalized expression values and expressed as z-scored delta activity relative to ISO. (B) Kaplan-Meier overall survival curves comparing the top 10% and bottom 10% Volumetric Mechanoadaptation Index subgroups in the TCGA SKCM cohort (n1=n2=46). Scatter plots show mean ± s.d. All RNA sequencing analyses were performed on adapted cells (5-day exposure) and compared to ISO.

We next asked whether patient tumors display a similar volumetric mechanoadaptive transcriptional axis. We derived a Volumetric Mechanoadaptation Index (VMI) from the adapted 50DI signature and scored TCGA SKCM tumors by transcriptional similarity to this state. In a UMAP projection of the cohort, VMI distributed as a continuous gradient with localized high-index regions (Extended Data Fig. 11B,C). Kaplan–Meier analysis showed worse overall survival for the highest VMI decile compared with the lowest decile (Fig. 6B). Together, these data connect the volumetric-history associated program to a patient-level transcriptional axis that is associated with poor outcome in human melanoma.

### Volumetric mechanoplasticity elevates immune visibility while shifting the glycocalyx toward reduced sialylation

The invasion-associated, drug-tolerant state raised a second therapy-relevant question: does volumetric mechanoplasticity also reshape tumor-intrinsic programs linked to immune visibility. We addressed this using adapted-state transcriptomics together with post-recovery protein readouts, and we then evaluated concordance in patient datasets. In the adapted transcriptome, IFN-γ response signatures were enriched in 50DI relative to ISO, whereas 25DI did not meet enrichment thresholds under the same analysis (Fig. 7A). This provides a direct rationale for immune visibility because IFN-γ programs are closely linked to induction of antigen processing and presentation pathways^60^. Consistent with this logic, antigen processing and presentation via MHC-I was enriched in 50DI relative to ISO, while the same gene set was not significant in 25DI (Fig. 7B). Adapted 50DI also increased transcripts encoding chemokines and components of the MHC-I antigen processing and presentation machinery (Fig. 7C). We next asked whether this signature persists beyond the adapted state. After return to isotonic conditions, primed 50DI cells displayed higher surface MHC-I by immunofluorescence, whereas primed 25DI remained near the ISO baseline (Fig. 7D,E). In TCGA SKCM, higher expression of chemokine and MHC-I machinery modules was associated with improved overall survival, with gene-level associations directionally consistent with an immune-engaged state (Extended Data Fig. 12A–C).

**Figure 7.**
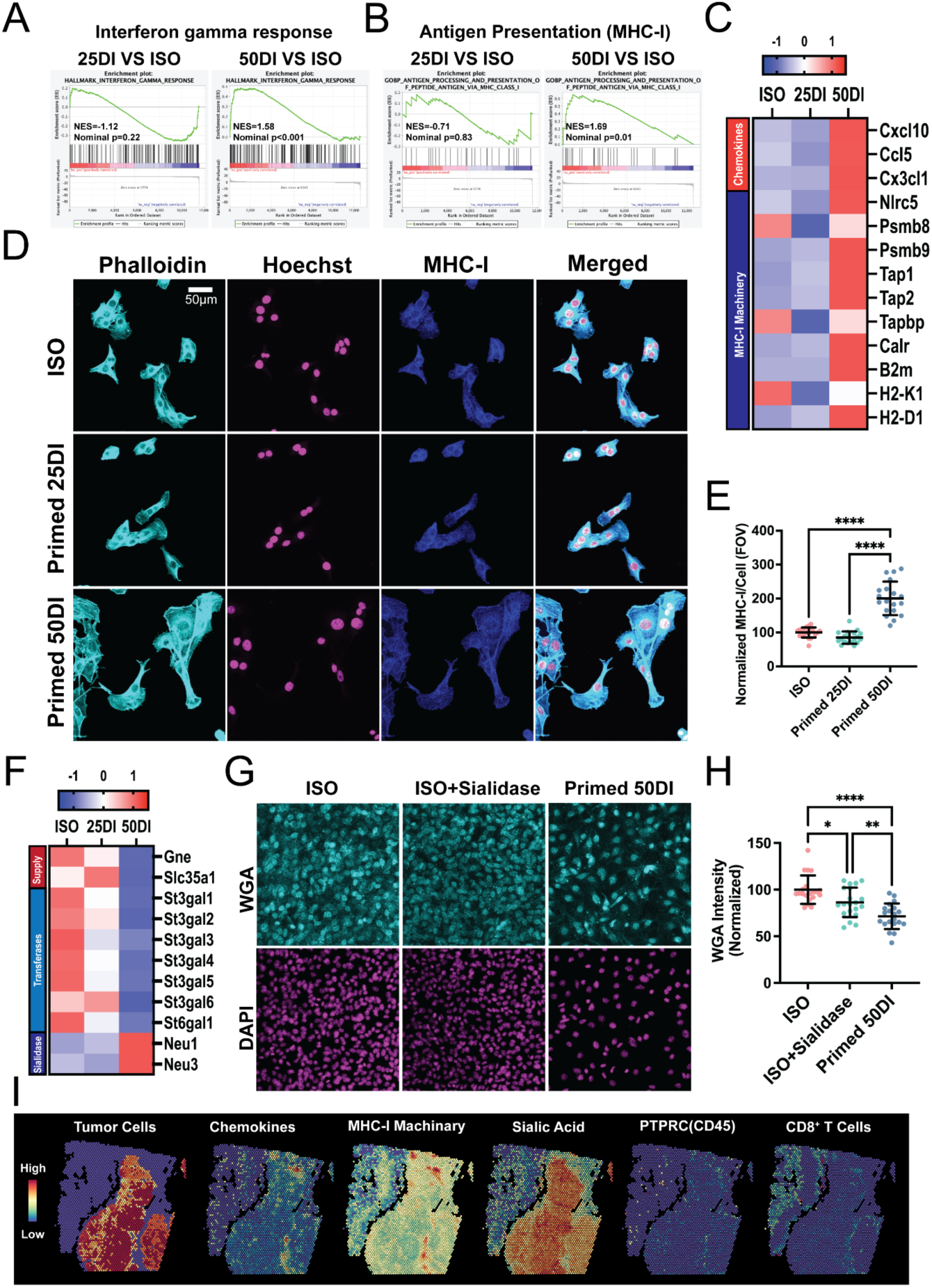
Volumetric mechanoplasticity increases antigen presentation and dampens surface sialylation in melanoma. (A) GSEA for the Hallmark interferon gamma response in adapted 25DI (top) and adapted 50DI (bottom) relative to ISO. (B) GSEA for antigen processing and presentation via MHC-I in adapted 50DI relative to ISO. The same gene set did not reach significance in adapted 25DI relative to ISO. (C) Heatmap of chemokines and MHC-I antigen processing and presentation machinery across ISO, adapted 25DI, and adapted 50DI from RNA sequencing. Heatmap colors show z-scored expression. (D) Post-recovery immunofluorescence images of ISO, primed 25DI, and primed 50DI cells stained for F-actin (phalloidin), nuclei (Hoechst), and surface MHC-I, with merged overlays shown below. (E) FOV quantification of normalized surface MHC-I intensity per cell across ISO, primed 25DI, and primed 50DI (n=19 cells per condition from 2 independent experiments). (F) Heatmap of sialic acid pathway genes across ISO, adapted 25DI, and adapted 50DI from RNA sequencing, grouped into sialic acid supply and transport, sialyltransferases, and sialidases. Heatmap colors show z-scored expression. (G) WGA lectin staining of ISO, ISO treated with exogenous sialidase, and primed 50DI cells, with DAPI shown below each condition. (H) FOV quantification of normalized WGA intensity across conditions (n=20 fields of view per condition from 4 independent experiments). (I) Spatial transcriptomics of a human melanoma section showing inferred tumor cell signal, CD8⁺ T cell signal, chemokine module score, MHC-I machinery module score, and a sialic acid supply and transferase module score derived from the genes in panel F. Scatter plots show mean ± s.d. All RNA sequencing analyses were performed on adapted cells (5 day exposure) and compared to ISO.

We next examined glycosylation programs that modulate immune recognition. In TCGA SKCM, a sialic acid supply and transferase module, comprising genes supporting sialic acid production and transfer to glycans, was higher in tumors than in non-tumor controls and this elevation was preserved across major melanoma genomic subtypes (Extended Data Fig. 12D,E). This pattern is consistent with broadly increased sialylation programs in melanoma, and sialylated glycans can function as suppressive glyco-checkpoints through inhibitory Siglec receptor engagement on immune cells^61–64^. Against this backdrop, adapted 50DI shifted the pathway in the opposite direction: sialic acid supply and transfer genes decreased, whereas sialidase genes increased relative to ISO and 25DI (Fig. 7F). As a phenotypic readout after recovery, primed 50DI showed reduced WGA lectin staining (Fig. 7G,H), matching the direction of an exogenous sialidase control that reduces surface sialic acids and can enhance tumor immunogenicity in multiple models^61–64^. Consistent with this interpretation, WGA binding in this context is sensitive to surface sialylation, as enzymatic removal of terminal sialic acids reduces WGA-dependent signal^65^.

Spatial analyses provided orthogonal context for these programs. In spatial transcriptomics, we visualized tumor-enriched signal together with a pan-immune marker readout based on PTPRC (CD45), inferred CD8⁺ T cell signal, and spatial module scores for chemokines, MHC-I machinery, and sialic acid supply and transfer within the same tissue section (Fig. 7I). Regions enriched for CD45 and CD8⁺ T cell signal overlapped and coincided with higher chemokine and MHC-I module scores. In contrast, the sialic acid module score was elevated in immune-sparse regions with low CD45 and low CD8⁺ T cell signal. Together, these multilayer data support a post-recovery state linked to volumetric mechanoplasticity that combines increased antigen presentation capacity with a coordinated shift of the sialic acid axis toward reduced sialylation-associated surface glycan display.

## Discussion

We define a history-dependent hydromechanical route to melanoma plasticity that we term volumetric mechanoplasticity. Using extracellular dilution as a controllable experimental handle, we compared two priming regimes that share sustained hypotonic exposure but differ in the magnitude and persistence of cell volume expansion during the priming window. This design helps distinguish effects associated with hypotonic stress from those linked to sustained volumetric enlargement and remodeling as drivers of post-recovery state changes. Across imaging, transcriptomic, functional, and in vivo readouts, the volumetric history biases melanoma toward a slow-cycling, invasion-competent program with greater tolerance to cytotoxic stress, while also increasing tumor-intrinsic immune visibility to CD8⁺ T cells. The central implication is therefore not simply that volume expansion alters phenotype, but that hydromechanical history can shift melanoma along an axis in which cytotoxic stress tolerance and immune visibility vary as separable dimensions of adaptation. In this framework, volumetric mechanoplasticity aligns drug tolerance with a concurrent vulnerability that immunotherapy can exploit.

A central feature of this framework is that the relevant input is history, not an acute hypotonic exposure. We explicitly stage acute exposure, multiday adaptation, and a post-recovery readout to distinguish immediate osmotic responses from phenotypes that persist after return to isotonic conditions^20,21,66^. The matched 25DI and 50DI comparison then serves as a stringent filter: phenotypes that are weak or absent in 25DI but prominent in 50DI are most consistent with dependence on robust volumetric expansion during priming rather than hypotonic stress alone. This logic reframes our findings as a history-encoded volumetric state that remains detectable after return to isotonic medium, rather than a transient response to hypotonic exposure. In this view, volumetric mechanoplasticity reflects a state-shift process, in which a defined hydromechanical experience biases cells toward a region of phenotypic space that remains occupied over the early post-recovery window, when treatment decisions and immune interactions can be decisive.

Our data implicate the nucleus as a plausible site where volumetric-history information could be integrated and retained, and they motivate a working model with explicit causal ordering. We propose that sustained volumetric expansion during priming increases cellular stress burden while remodeling nuclear envelope composition and chromatin organization. The lamina shift together with persistent post-recovery chromatin decondensation suggests that changes at the nuclear periphery and in chromatin compaction may contribute to storing aspects of the history as an altered nuclear mechanical context^36^. In parallel, stress signaling and checkpoint engagement, including p53-linked programs, could provide a route by which this stored nuclear state is translated into durable suppression of DNA replication and mitotic programs^67^. We do not claim a fully resolved molecular chain; rather, the observations support a coherent sequence in which volumetric expansion during priming establishes a persistent alteration in nuclear architecture^36^, this alteration biases stress and checkpoint set points^40^, and the resulting checkpoint tone limits proliferative capacity even after return to isotonic conditions^67^. Making the model explicit is valuable because it generates falsifiable predictions. If nuclear remodeling is required for persistence, then perturbations that prevent lamin remodeling or restore chromatin compaction should weaken the post-recovery slow-cycling phenotype. Conversely, inducing comparable nuclear remodeling in isotonic conditions, without hypotonic priming, may recapitulate key features of the primed state.

Volumetric mechanoplasticity reframes the proliferation–invasion tradeoff as an output of hydromechanical history rather than solely of canonical transcriptional switching. The volumetric regime engages EMT- and adhesion-related remodeling, shifts the integrin expression repertoire toward fibronectin-associated programs, and is accompanied by reinforced F-actin architecture together with increased motility in three-dimensional collagen matrices. In melanoma, where transitions between growth-biased and invasion-biased programs are central to therapeutic escape, these shifts have direct implications for residual disease and dissemination^1^. Our data therefore support the interpretation that sustained volumetric expansion history can act as a physical bias that alters which plasticity trajectories are preferentially occupied. More broadly, this view is consistent with evidence that multiday mechanical histories can prime durable state shifts across systems, including melanoma^20,21,66^. Importantly, this bias is compatible with slower net population expansion. Reduced divisions per unit time does not necessarily imply fewer surviving cells under treatment if stress tolerance and dispersal are simultaneously altered.

The cytotoxic-therapy tolerance program illustrates this point most directly. Volumetrically primed cells engage antioxidant and glutathione programs, anti-ferroptosis defenses, and survival signaling, consistent with increased capacity to buffer oxidative and genotoxic stress^55,58^. This apparent paradox is clinically relevant because treatment outcome is driven by attrition of viable tumor cells, not by proliferation rate alone^68^. A slow-cycling population can dominate the residual pool if it experiences reduced therapy-mediated loss^69^. In addition, dissemination can increase residual burden by redistributing viable cells into microenvironmental niches that offer protection^70^. In this light, volumetric mechanoplasticity provides a concrete route by which physical microenvironment history can generate a relapse-enabling combination of traits: survival under cytotoxic stress together with spatial escape.

The most unexpected output of volumetric mechanoplasticity is the immune visibility phenotype. Volumetric history induces interferon-related programs, increases antigen processing and presentation machinery, elevates surface MHC-I after recovery, and is accompanied by a coordinated shift in the sialylation axis, including reduced sialylation-associated modules and reduced WGA lectin binding. Two clarifications are important. First, we do not interpret this as a general rule that stress adaptation enhances immune recognition; we present it as a context-specific co-occurrence within this history-defined state. Second, this immune visibility output strengthens the core conceptual claim: volumetric mechanoplasticity can generate a state that is more tolerant to cytotoxic therapies yet comparatively accessible to cytotoxic lymphocytes. Several mechanistic routes are plausible and directly testable. Chromatin decondensation together with stress signaling could elevate basal inflammatory tone^67^ and increase antigen presentation capacity^60^, whereas reduced sialylation could diminish an inhibitory glycan layer that limits immune synapse formation and engages Siglec-mediated suppression^61–64^. Practically, these results suggest that residual, stress-adapted melanoma states may not be uniformly immune silent.

Patient relevance is supported by the Volumetric Mechanoadaptation Index (VMI), a transcriptional similarity score derived from the adapted 50DI signature, which identifies TCGA SKCM tumors in which higher VMI is associated with poorer overall survival. These associations do not establish causality and are vulnerable to confounding inherent to bulk data, including variable immune infiltration that can influence immune-related modules. Nonetheless, the signal is unlikely to be purely artifactual. When considered alongside the spatial neighborhood structure of nuclear size observed in human melanoma tissue, our data support the idea that melanoma ecosystems contain spatially structured subpopulations that may be consistent with distinct hydromechanical histories (Fig. 1A,B; Extended Data Fig. 1). The priority now is to move from association to linkage. Spatially resolved measurements should test whether enlarged, stress-marked tumor regions also show increased antigen presentation and altered glycosylation signatures, and whether these regions emerge or expand under therapy.

Several limitations sharpen the scope and strengthen the interpretation. First, we use extracellular osmolarity as a controllable experimental handle to impose defined hydromechanical histories and to distinguish hypotonic stress from sustained volume expansion as much as possible. This is not a claim that tumors are chronically hypotonic. The in vivo plausibility we invoke is the existence of hydromechanical niches that can bias water flux and volume regulation over time, including edematous and glycosaminoglycan-rich matrices that retain fluid and reshape transport^9,12^. Consistent with this framing, swollen subpopulations and gradients in cell and nuclear size have been reported in 3D tumor models and in patient specimens^15–17^. Second, osmotic downshift and swelling magnitude covary and cannot be perfectly decoupled. As a result, the 50DI condition could reflect a threshold response to combined perturbations rather than an effect attributable to volume expansion alone. Our design reduces, but does not eliminate, this ambiguity, which is why we interpret 25DI as a conservative comparator rather than as definitive proof of a single causal variable. In practice, direct and isolated control of multiday volume expansion remains difficult, and hypotonic dilution has therefore remained a tractable lever in related works^14,15,17,18,32^. Third, clinical association analyses are constrained by bulk data and historical context. TCGA signals mix tumor-intrinsic state with immune composition and derive from a treatment era that differs from current practice. Accordingly, we do not infer that VMI predicts modern immunotherapy response; rather, the association indicates that the state is detectable at scale in patient data and warrants targeted investigation in contemporary, clinically annotated cohorts.

Within these boundaries, volumetric mechanoplasticity provides a mechanistic lens for a clinical observation that is frequently reported in melanoma: tumors can progress on targeted therapy yet remain responsive to immune checkpoint blockade or adoptive cell therapy in a subset of patients^5–8^. One plausible scenario is that residual cells occupying hydromechanical niches that bias sustained volumetric expansion acquire stress-adapted survival programs without necessarily acquiring the immune concealment phenotypes that would limit cytotoxic lymphocyte recognition. In this framing, volumetric mechanoplasticity is not presented as a clinical sequence. Rather, it clarifies how drug tolerance can emerge without mandatory immune escape and it motivates actionable hypotheses. If volumetric history increases immune visibility while installing cytotoxic tolerance, then combining cytotoxic therapies with immune strategies that amplify recognition and immune synapse function may be most effective during early residual disease windows, a direction that is consistent with emerging combination approaches in clinical trials^71^. Conversely, if the same history elevates MHC-I and reduces sialylation-associated programs, then glycan checkpoint targeting and interferon-axis modulation become rational interventions to widen the gap between immune control and drug tolerance. More broadly, hydromechanical history represents a complementary layer of cellular memory that may help explain which residual states persist under therapy alongside transcriptional and metabolic history.

## Materials and Methods

### Ethics, approvals, and study oversight

Human melanoma histology analyses used a deidentified melanoma tissue microarray containing biopsy cores provided by the Yale Department of Pathology. Deidentification was completed by the provider prior to transfer.

### Cell lines, culture conditions, and volumetric priming

B16F0 and B16F10 mouse melanoma cells were obtained from ATCC. YUMM and YUMMER1.7 cells were provided by Marcus Bosenberg’s laboratory at Yale. Cells were maintained at 37 °C with 5% CO₂ in DMEM for B16 lines or DMEM F12 for YUMM and YUMMER1.7, supplemented with 10% fetal bovine serum and 1% penicillin streptomycin. Cells were passaged every 2 to 3 days. Isotonic control medium (ISO) refers to the complete growth medium described above. Hypotonic media were generated by diluting complete growth medium with deionized water to final fractions of 25% (25DI) or 50% (50DI) deionized water. To match total nutrient delivery across conditions, the total culture volume per well was increased in proportion to dilution for 25DI and 50DI so that the total amount of base medium per well matched ISO. Medium osmolality was measured using a vapor pressure osmometer (Vapro 5600, EliTech).

Hydromechanical history was operationalized using three temporal states. The acute state reflects a 2 h exposure to ISO, 25DI, or 50DI. The adapted state reflects continuous culture in 25DI or 50DI for 5 days, with ISO cultured in parallel. The primed state reflects return to ISO after adaptation, followed by reseeding and analysis 24 h later. Adaptation was initiated by seeding cells in ISO and allowing attachment overnight, followed by replacement with 25DI or 50DI to start the 5 day adaptation window. Cultures were started at approximately 1% confluence for ISO, approximately 2% confluence for 25DI, and approximately 10% confluence for 50DI to prevent overconfluence by day 5 while preserving sufficient cell yield. Media were refreshed every 2 to 3 days. After 5 days, cells were trypsinized, counted, reseeded into ISO at assay specific densities, and analyzed after 24 h recovery to define the primed state.

### Microscopy, staining workflows, enzyme treatment, and general image quantification

Brightfield imaging was performed on an inverted microscope (Leica DMi1). Confocal imaging was performed on a Leica SP8 series confocal microscope or a Nikon FV1000 microscope. Objectives of 10×, 20×, 40×, or 60× were used depending on the assay. Within each experiment, acquisition settings were held constant across conditions to enable quantitative comparisons. Unless stated otherwise, fluorescence z stacks spanning 10 μm were acquired and quantified using maximum intensity projections.

For fixed cell staining, cells were fixed in 4% paraformaldehyde in PBS for 15 min, washed in PBS, and blocked in 3% bovine serum albumin in PBS for 1 h at room temperature. For surface MHC-I staining, primed cells were replated in ISO for post recovery analysis, fixed in 4% paraformaldehyde in PBS for 10 min, and blocked in 3% BSA in PBS for 1 h without permeabilization. Samples were incubated with an antibody recognizing mouse MHC-I (Thermal Scientific, 12-5998-82) at 1:20 dilution for 30 minutes at room temperature. Nuclei were stained with Hoechst 33342 or DAPI for 15 min at 1:1000 dilution. F-actin was stained using rhodamine phalloidin (Invitrogen) at 1:2000 for 30 min. Image processing and quantification were performed in Fiji using custom scripts. When single cell quantification was performed, segmentation masks were generated manually or through Analyze Particles FIJI plug-in and all analysis parameters were held constant across conditions within each experiment.

Surface glycan changes were assessed by staining cells with wheat germ agglutinin lectin (WGA 647, Thermo Fisher, W32466). Cells were incubated with WGA at 1 μg/ml for 10 min at room temperature, washed three times in PBS, and imaged by fluorescence microscopy. WGA intensity was quantified using whole cell masks derived from cell GFP and normalized to ISO within each experiment. For desialylation controls, ISO cells were treated with sialidase (Sigma, N2876) at 1 U/ml for 1 h at 37 °C in complete medium, washed, and processed for WGA staining in parallel.

### Acute swelling assays and 3D volume reconstruction

Acute volumetric response assays were performed by exposing cells for 2 h to ISO, 25DI, or 50DI. For 3D volume reconstruction, z stacks were acquired with a z step of 1 μm and single cell volume was reconstructed in Imaris using surface based segmentation with the GFP channel as the primary signal. Sphericity was computed from reconstructed 3D surfaces using the standard Imaris implementation. Projected cell area was quantified from maximum intensity projections using Fiji and custom scripts with identical parameters across conditions. Confocal-based volume reconstruction has been validated against high resolution structured illumination microscopy measurements of cell volume^18^

### Cell and nuclear morphology, scaling coupling, and additional shape metrics

Projected nuclear area and projected cell area were quantified in primed cells from confocal images using segmentation masks generated in Fiji with fixed parameters within each experiment. Additional nuclear and cell shape descriptors, including major axis length, circularity, and aspect ratio, were computed from the same masks.

### Chromatin condensation index in primed cells

Chromatin state in primed cells was quantified from Hoechst or DAPI images using an edge texture based chromatin condensation index adapted from prior image based chromatin packing measurements^37,38^. Briefly, projected nuclear images were intensity normalized in Fiji. Nuclei were segmented to generate masks. Edge maps were generated using a Sobel edge filter (Fiji, Find Edges) and skeletonized to produce single pixel edge networks (Fiji, Skeletonize). The chromatin condensation index was computed per nucleus as skeleton area divided by nuclear cross-sectional area.

### Population expansion during adaptation and post recovery regrowth

Population expansion during adaptation was quantified by imaging parallel cultures in ISO, 25DI, and 50DI at day 1, day 3, and day 5 through capturing random FOVs. Cell number was quantified by manual counting in Fiji from multiple nonoverlapping fields per condition and time point. Growth was reported as fold change relative to each condition’s baseline within the same experiment, and replicate level log_10_(day 5 divided by baseline) values were used for statistics. Post recovery regrowth was quantified by replating ISO controls and adapted cells into ISO 3D collagen matrix and imaging a 100μm Z range over 36 h. Cancer cell max z-projection masks were generated and were used for segmentation. Thresholding and morphological cleanup were applied identically across conditions. Percent area covered by cancer cells was computed for each time point, and endpoint area at 36 h was summarized across images and independent experiments.

### F-actin ridge analysis and cytoskeletal quantification

Actin organization was quantified in primed cells after phalloidin staining and confocal imaging. Single cell regions were segmented manually. Mean phalloidin fluorescence intensity was computed per cell. Ridge-like F-actin structures were extracted by custom Fiji script using Ridge Detection plugin. Ridge number per cell was defined as the count of disconnected skeleton components in each of the z slides, and total ridge length per cell was defined as the summed skeleton length in all z slides. All analysis parameters were held constant within an experiment.

### 3D collagen and hyaluronan collagen motility assays

Three-dimensional matrices were prepared using acetic acid solubilized type I rat tail collagen (Corning). Collagen, 10× PBS, double distilled water, and 0.5 N NaOH were mixed on ice and neutralized to reach a final collagen concentration of 1 mg/ml. Single cell suspensions were mixed with collagen, transferred into 96 well plates kept on ice, polymerized at 37 °C, and overlaid with ISO medium for time-lapse imaging. For collagen plus hyaluronan matrices, hyaluronan (Biosynth, FH63426, extra low) was added before polymerization to reach a final concentration of 1.2 mg/ml following an established protocol^18,72^. Time lapse imaging was performed with a frame interval of 1 hour. Tracks were obtained using Fiji TrackMate. Cell morphology metrics including aspect ratio, circularity, and solidity were computed from binary masks. Velocity and mean squared displacement were computed from trajectories using Fiji and MATLAB.

### RNA sequencing and computational analyses

RNA sequencing was performed on ISO, adapted 25DI, and adapted 50DI B16F0 cells using two independent experiments with separate cultures and RNA extractions. Total RNA was extracted using the RNeasy Micro Kit (Qiagen) and sequencing was performed by the Yale Center for Genome Analysis. FASTQ reads were aligned to the mouse genome (GRCm38) using STAR (2.7.9a) in 2 pass mode. Alignments were indexed using samtools (1.11) and gene level counts were generated using HTSeq (0.13.5). Count matrices were analyzed in R (v4.1.3). Differential expression was performed using DESeq2 with significance defined using adjusted P value thresholds as indicated in figure panels. Hallmark pathway activity was quantified using GSVA^33^ on the MSigDB Hallmark gene set collection^34^. ΔGSVA scores were computed relative to ISO using the ISO mean across replicates as baseline. For ranked enrichment analysis, genes were ordered by signed differential expression statistics and analyzed by GSEA Preranked^35^, with leading edge genes extracted for visualization when indicated. EnrichmentMap networks were visualized in Cytoscape using EnrichmentMap, where nodes represent enriched terms and edges represent gene overlap similarity.

### Patient level analyses, VMI derivation, and public spatial transcriptomics

TCGA SKCM expression and survival analyses were performed using GEPIA2^73^, using publicly available TCGA SKCM data. A Volumetric Mechanoadaptation Index (VMI) was derived from similarity to the adapted 50DI transcriptional program, with gene lists and scoring details provided in Source Data. For survival analysis, patients were stratified into the top 10% and bottom 10% VMI groups using cohort specific cutoffs and overall survival was compared using Kaplan–Meier analysis with log-rank testing.

Public human spatial transcriptomics data were analyzed using the SpatialTME portal^74^ and a publicly released 10x Genomics Visium melanoma dataset displayed within SpatialTME (dataset: Melanoma_IFStainedFFPE_CytAssist_10x, 10x Visium Datasets). Tumor cell and CD8 T cell signals were inferred using marker based scoring implemented in the portal. Module scores were computed using the gene lists defined in figure panels, and spatial heatmaps were exported from the portal for visualization.

### Human melanoma histology: nuclear segmentation, heatmaps, and spatial statistics

Human melanoma histology was analyzed from a representative H&E section imaged as a tiled dataset on an EVOS M7000 Imaging System (Invitrogen). Nuclear masks were generated using a Cellpose SAM workflow^25^. Nuclear area was computed per nucleus and normalized to the mean nuclear area across the section. Heatmaps were generated by mapping segmented nuclei to spatial coordinates and displaying log₂(nuclear area divided by mean nuclear area).

Spatial autocorrelation of nuclear area was quantified using both global and local Moran statistics^30,31^. Moran statistics were computed on nuclear area values using inverse distance weighted k nearest neighbor spatial weights (k = 24). Global Moran’s I significance was assessed by permutation, randomly permuting nuclear area across fixed spatial coordinates (999 permutations) to obtain an empirical null distribution and a two sided permutation P value. Local indicators of spatial association were computed using the same spatial weights and permutation based local significance testing at alpha = 0.05 to classify nuclei into High-High and Low-Low clusters, High-Low and Low-High spatial outliers, or not significant nuclei.All spatial statistics and maps were computed on the same nuclei used for the corresponding main figure panels.

### Statistics, experimental design, replication, randomization

Imaging fields of view were acquired randomly to minimize selection bias. Statistical analyses were performed in R and GraphPad Prism 10. Two group comparisons used unpaired two sided t-tests. Multi-group comparisons used one way ANOVA with Tukey post-hoc. Exact statistical tests, sample sizes, and the number of independent experiments are reported in the figure legends. Data are shown as mean ± s.d. Statistical significance was annotated as *P < 0.05, **P < 0.01, ***P < 0.001, and ****P < 0.0001.

## Data and code availability

RNA sequencing data have been uploaded to the NCBI Sequence Read Archive under BioProject PRJNA955121. TCGA and other public datasets used are available from their respective portals. Quantitative source data supporting the figures are provided with the Article and Extended Data. Custom scripts are available from the authors upon reasonable request.

## Author contributions

X.Z. and M.M. designed research; X.Z., D.Z., X.G., A.P.Y., X.S., I.D., and M.M. performed research; X.Z., D.Z., M.M., L.L.M, R.K.J., analyzed data; and X.Z., M.M, L.L.M., R.K.J. wrote the paper.

## Acknowledgments

We acknowledge Dr. Gerald Shadel for helpful discussions. We acknowledge Dr. Rong Fan and Dr. Yanxiang Deng for their support with the bioinformatics analysis and access to the EVOSTM M7000 Imaging System (InvitrogenTM).

## Extended Data

**Extended Data Fig. 1.**
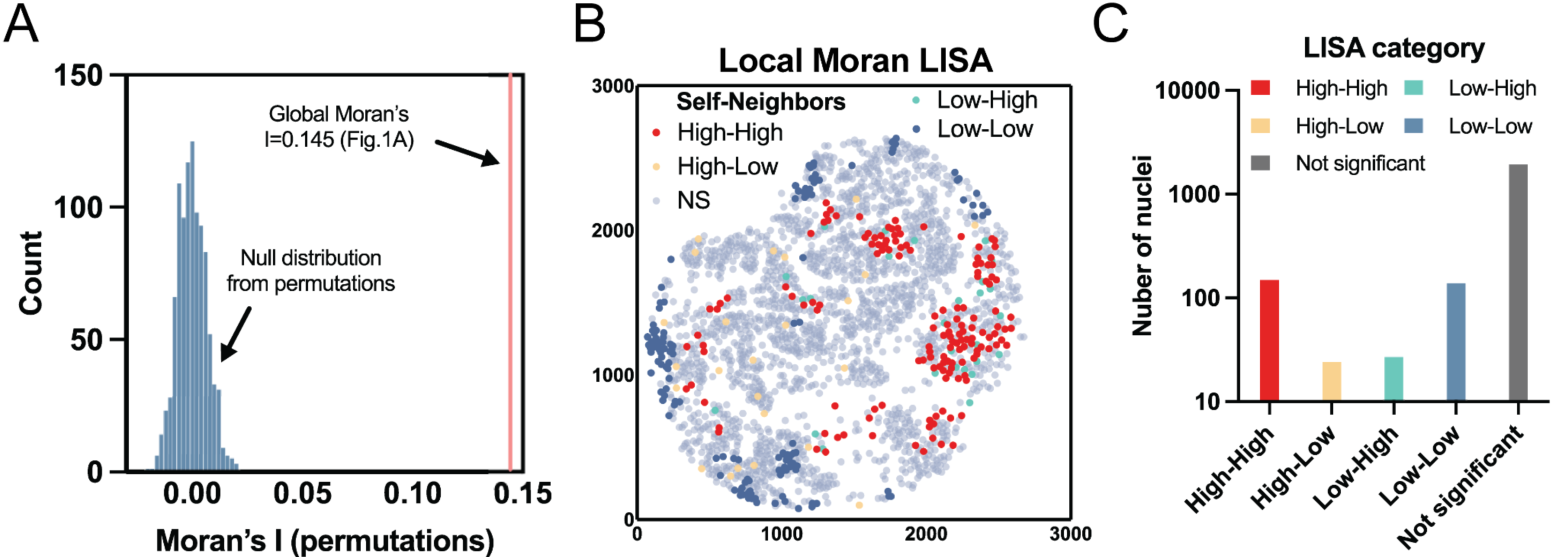
Spatial statistics and unsupervised segmentation confirm nonrandom nuclear size neighborhoods in human melanoma. (A) Global spatial autocorrelation of nuclear size. Global Moran’s I of the sample was computed on per nucleus nuclear area using inverse distance weighted k nearest neighbor spatial weights (k = 24), with Y coordinates flipped to match image orientation. The histogram shows the null distribution of Moran’s I generated by permuting nuclear area values across fixed nuclear coordinates (999 permutations), preserving tissue geometry while breaking any nuclear size neighborhood structure. The vertical line marks the observed Moran’s I (I = 0.1456), indicating significant positive spatial autocorrelation (two sided permutation p = 0.001). (B) Local spatial association map (LISA) for nuclear size neighborhoods. Local Moran statistics were computed using the same spatial weights (k = 24) to localize where clustering occurs within the section. Each nucleus was assigned a category using permutation based local significance testing at alpha = 0.05: High–High (large nucleus surrounded by large nuclei), Low–Low (small nucleus surrounded by small nuclei), High–Low (large nucleus surrounded by small nuclei), Low–High (small nucleus surrounded by large nuclei), or Not significant (no local spatial association at the chosen threshold). Colors correspond to the category key shown in the panel. (C) Summary of local spatial structure. Counts of nuclei in each LISA category from B, ordered as Not significant, Low–High, High–Low, Low–Low, High–High, summarizing the prevalence of significant clusters and spatial outliers across the tissue section. All analyses were performed on the same segmented nuclei used in Fig. 1A,B (n = 2261).

**Extended Data Fig. 2.**
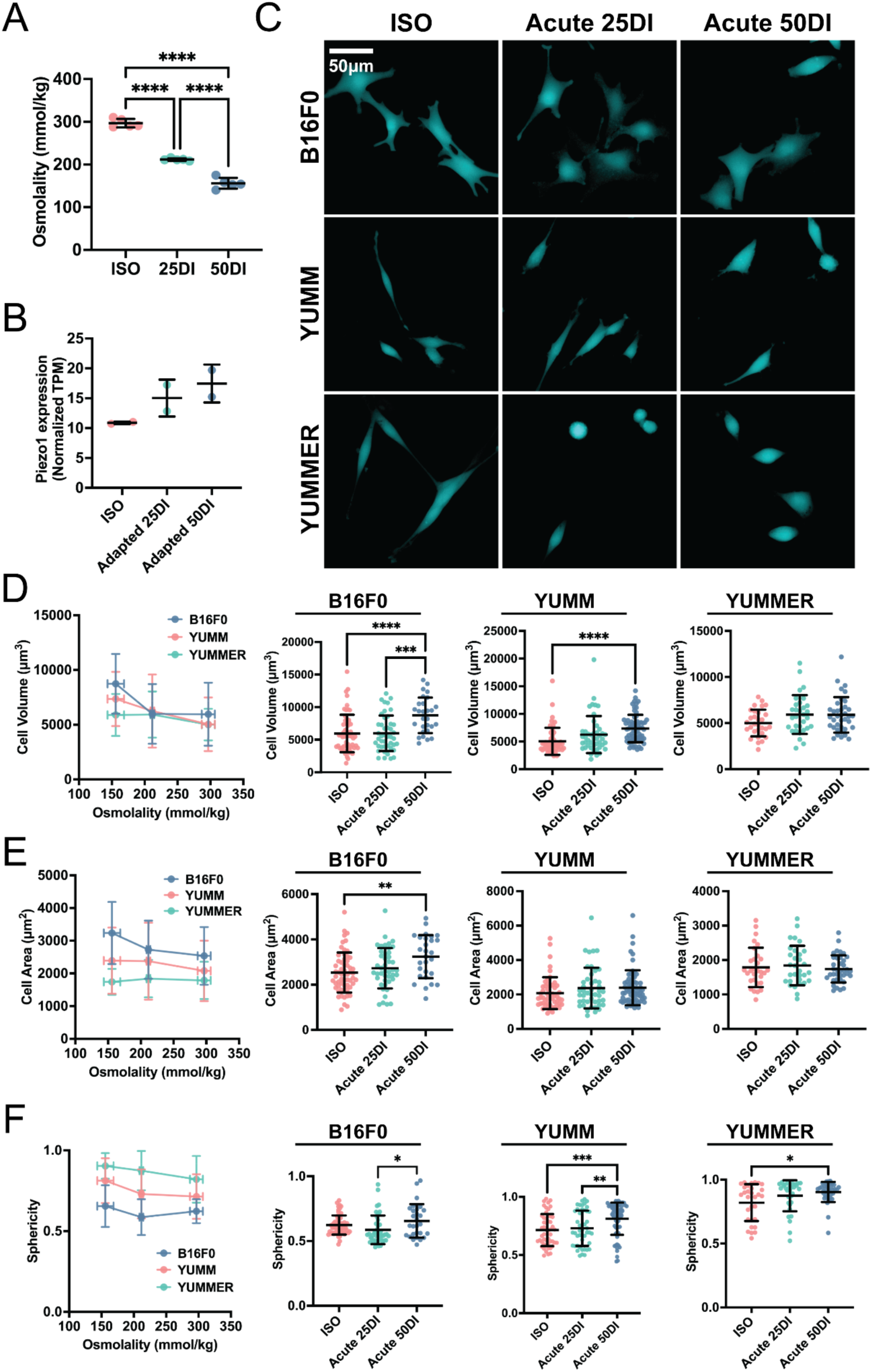
Calibration of osmotic inputs and acute swelling responses across melanoma backgrounds. (A) Medium osmolality across ISO, 25DI, and 50DI generated by DI water dilution (n = 5 technical measurements). (B) Piezo1 expression in adapted B16F0 cells across ISO, adapted 25DI, and adapted 50DI. (C) Representative fluorescence images of B16F0, YUMM, and YUMMER1.7 melanoma cells in ISO and during acute hypotonic exposure (2 h) across ISO, 25DI, and 50DI. Scale bars as indicated. (D–F) Acute single cell responses after 2 h across ISO, 25DI, and 50DI, quantified as cell volume (D), projected cell area (E), and sphericity (F) for B16F0 (n ≥ 28), YUMM (n ≥ 45), and YUMMER1.7 (n ≥ 30), from 2 independent experiments per line. These data confirm that 50DI induces robust swelling across melanoma backgrounds, while genetic background influences the accompanying shape response, including increased rounding in YUMM and YUMMER1.7. Scatter plots show mean ± s.d.

**Extended Data Fig. 3.**
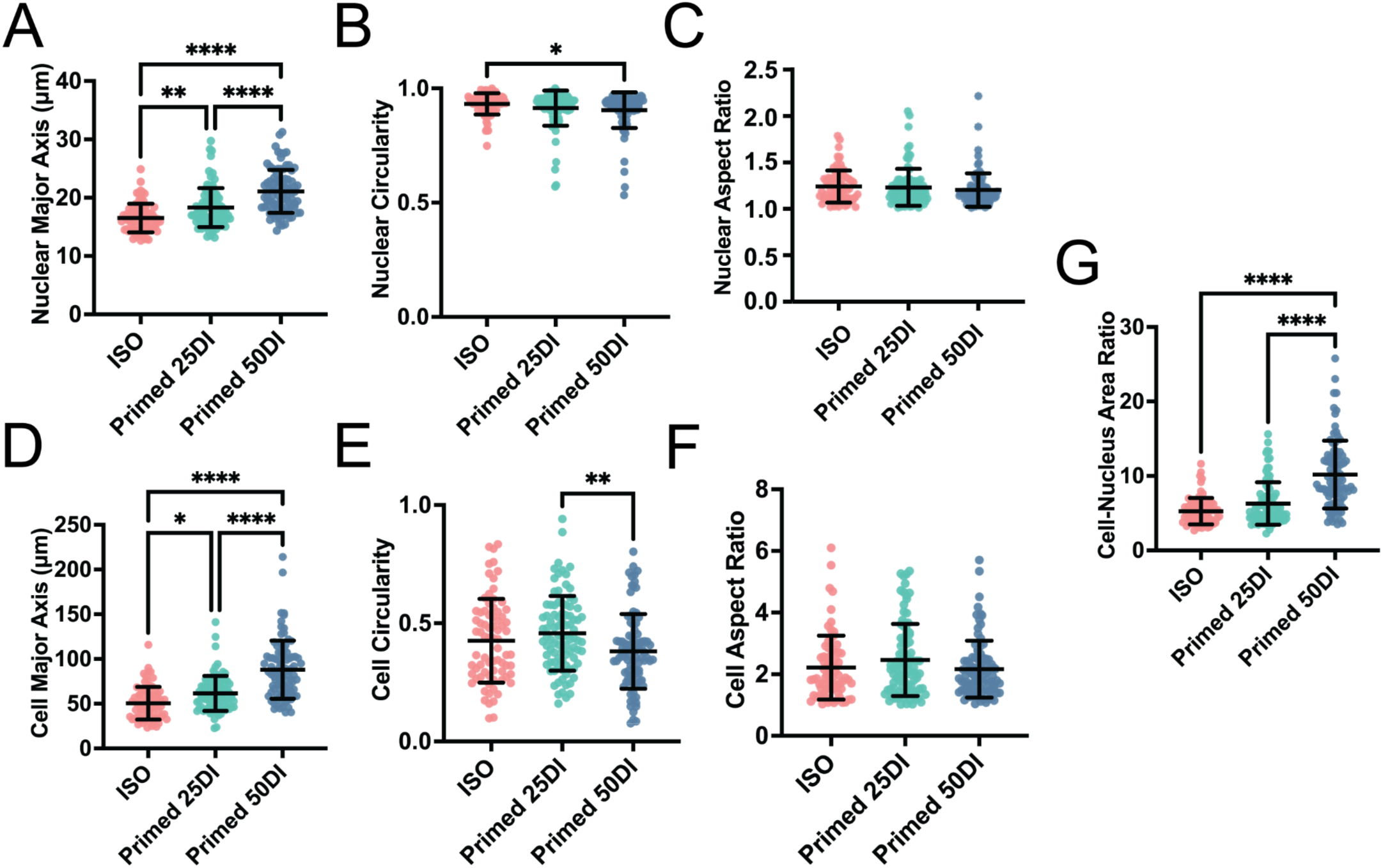
Additional nuclear and cell shape metrics after priming. (A–C) Single cell quantification of nuclear major axis (A), nuclear circularity (B), and nuclear aspect ratio (C) across ISO, primed 25DI, and primed 50DI. (D–F) Single cell quantification of cell major axis (D), cell circularity (E), and cell aspect ratio (F) across ISO, primed 25DI, and primed 50DI (n ≥ 76 cells per condition, 3 independent experiments). (G) Single cell quantification of the cell to nucleus projected area ratio across ISO, primed 25DI, and primed 50DI (n ≥ 76 cells per condition, 3 independent experiments). These metrics support the interpretation that primed 50DI produces a durable enlargement with modest distortion in shape descriptors relative to ISO, whereas primed 25DI produces smaller shifts. Scatter plots show mean ± s.d.

**Extended Data Fig. 4.**
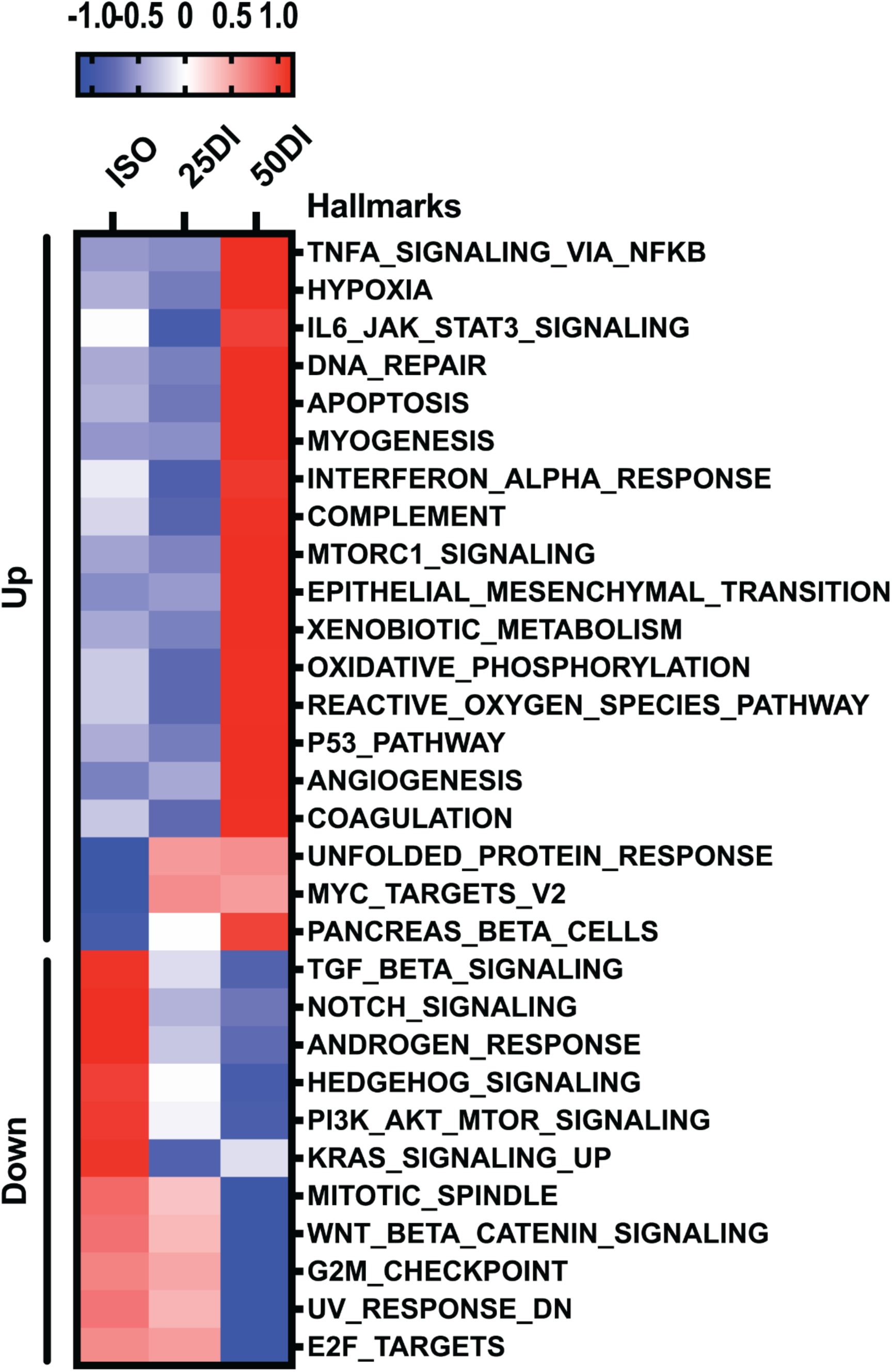
Hallmark GSVA heatmap across ISO, adapted 25DI, and adapted 50DI. Heatmap of selected MSigDB Hallmark gene sets scored by GSVA across ISO, adapted 25DI, and adapted 50DI. Values are displayed as z scored enrichment across samples. Gene sets are grouped by direction and by whether the dominant shift is shared across 25DI and 50DI or enriched in 50DI. All RNA sequencing analyses were performed on adapted cells (5 day exposure) and compared to ISO.

**Extended Data Fig. 5.**
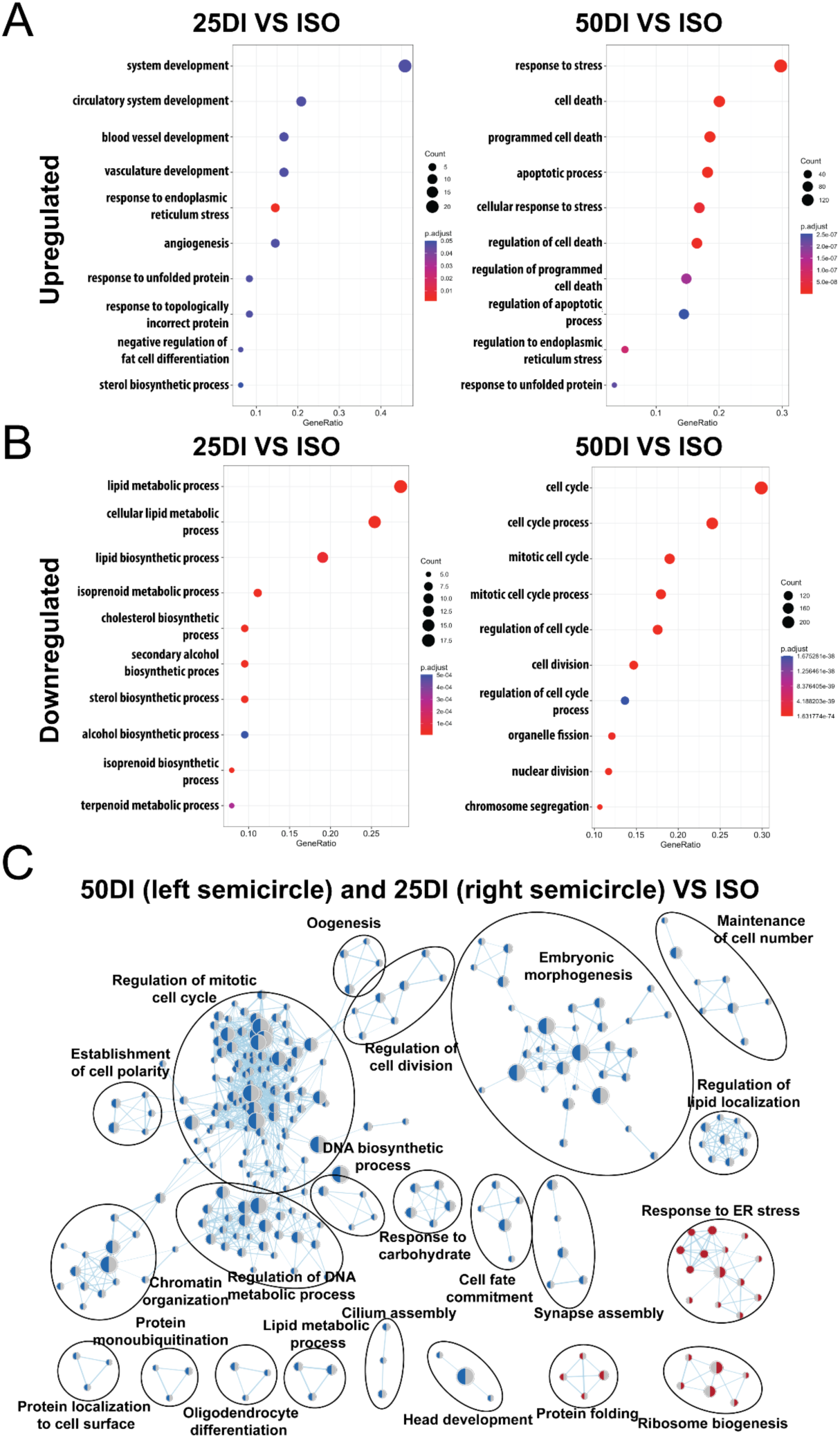
Gene Ontology enrichment and network view resolve a shared stress core and 50DI selective suppression of proliferative control. (A) Gene Ontology biological process enrichment for genes upregulated in adapted 25DI relative to ISO and in adapted 50DI relative to ISO. Dot size indicates gene count and color indicates adjusted P value. Terms are displayed for each comparison with redundant terms filtered for clarity. (B) Gene Ontology biological process enrichment for genes downregulated in adapted 25DI relative to ISO and in adapted 50DI relative to ISO, highlighting selective suppression of cell cycle, DNA replication, and mitotic programs in 50DI. Dot size indicates gene count and color indicates adjusted P value. (C) EnrichmentMap network visualization of enriched Gene Ontology biological processes from panels A and B. Nodes represent enriched terms and edges represent gene overlap similarity between terms. Node color encodes enrichment direction and condition association as defined in the panel. This analysis recapitulates a shared ER stress and protein quality control neighborhood and a large proliferative control neighborhood selectively suppressed in adapted 50DI. All RNA sequencing analyses were performed on adapted cells (5 day exposure) and compared to ISO.

**Extended Data Fig. 6.**
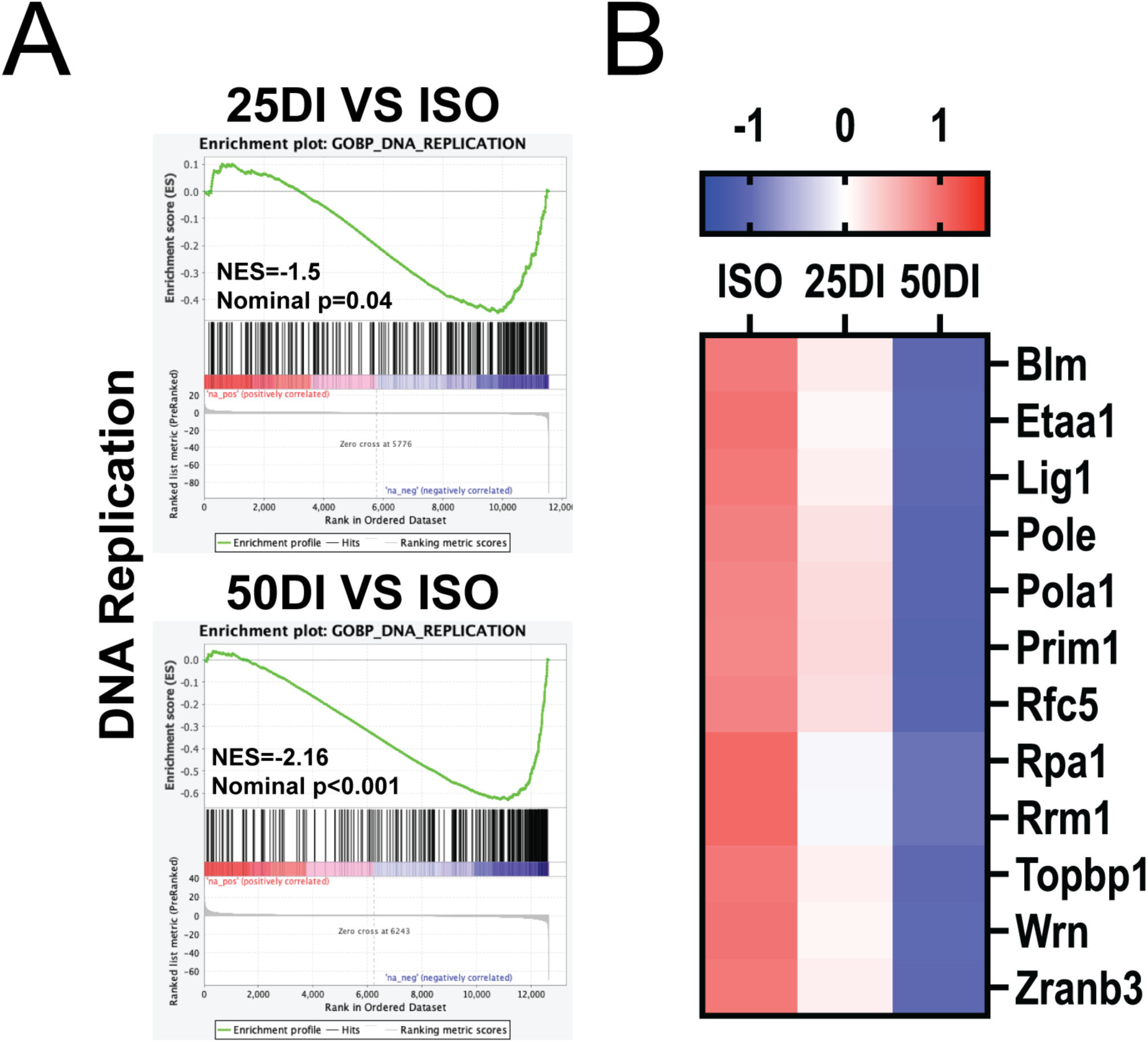
Replication associated gene sets are selectively suppressed in adapted 50DI. (A) GSEA for a curated DNA replication gene set comparing adapted 25DI versus ISO and adapted 50DI versus ISO, using the ranked gene list from differential expression analysis. (B) Heatmap of core DNA replication genes across ISO, adapted 25DI, and adapted 50DI, displayed as z scored expression across samples. All RNA sequencing analyses were performed on adapted cells (5 day exposure) and compared to ISO.

**Extended Data Fig. 7.**
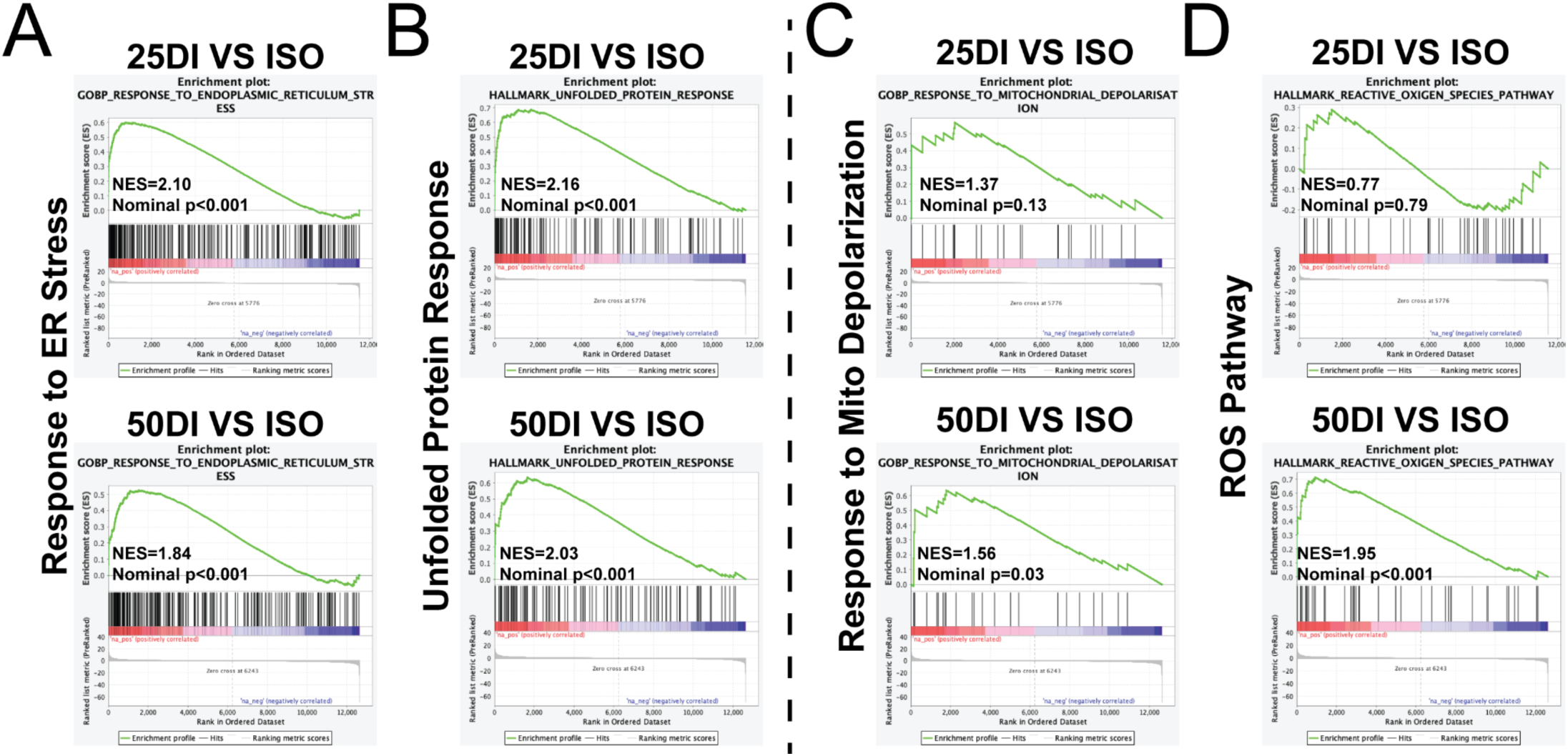
Pathway level enrichment and acute ROS imaging support stronger multi axis stress in 50DI. (A–D) GSEA for response to ER stress (A), unfolded protein response (B), mitochondrial depolarization (C), and ROS pathway (D) in adapted 25DI (upper) and adapted 50DI (lower) relative to ISO. All RNA sequencing analyses were performed on adapted cells (5 day exposure) and compared to ISO.

**Extended Data Fig. 8.**
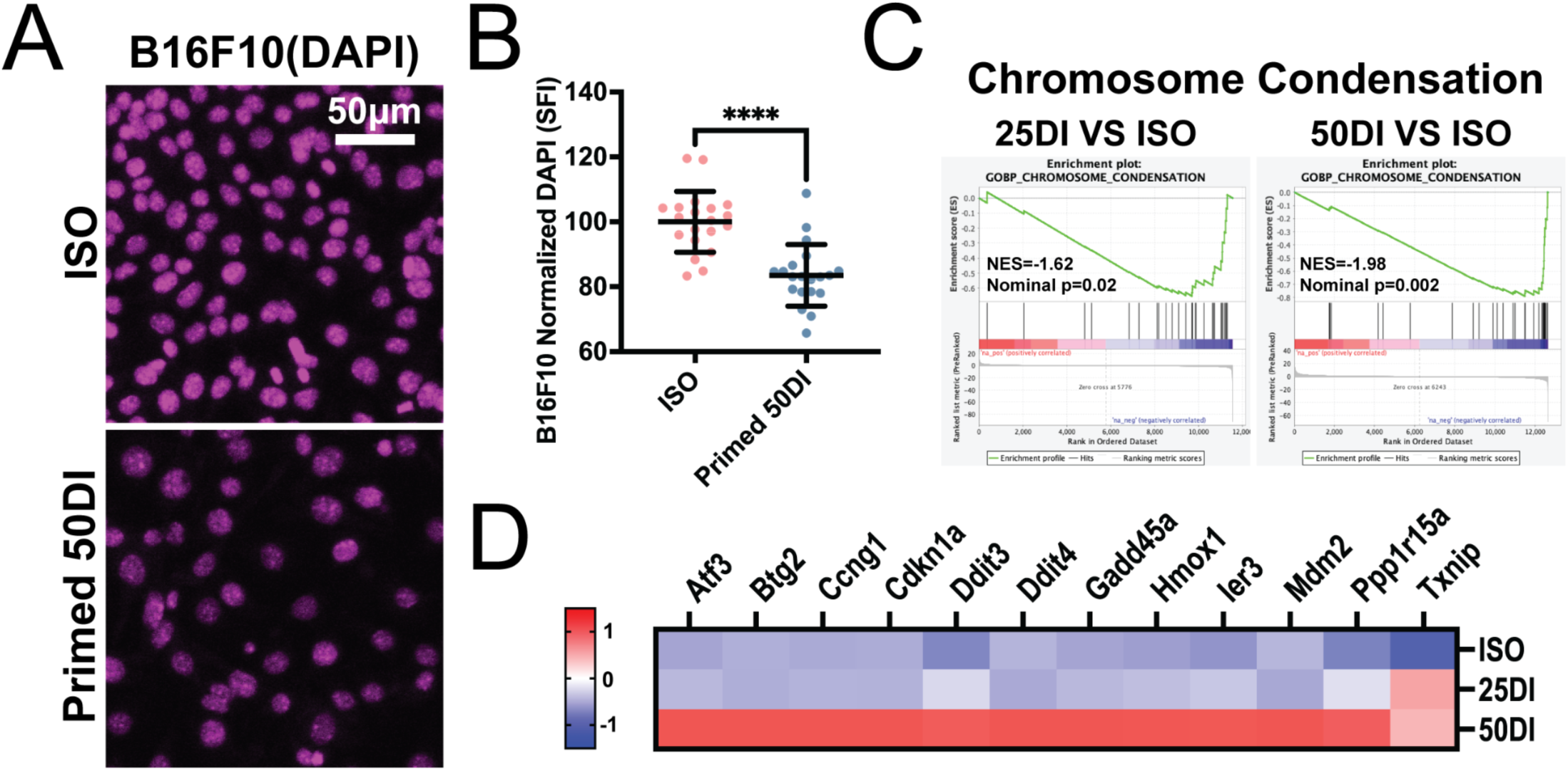
Cross line nuclear readouts and stress target genes support a volumetric primed state. (A) Representative nuclear staining images of primed B16F10 cells stained with DAPI across ISO and primed 50DI. Scale bar as indicated. (B) Single nucleus quantification of normalized DAPI intensity in primed B16F10 nuclei (n ≥ 20 per condition from 4 independent experiments). (C) GSEA for a chromosome condensation gene set comparing adapted 25DI versus ISO and adapted 50DI versus ISO in B16F0. (D) Heatmap of selected stress and p53 related target genes across ISO, adapted 25DI, and adapted 50DI from B16F0 RNA sequencing. Values are displayed as z scored expression across samples. Scatter plots show mean ± s.d. All RNA sequencing analyses were performed on adapted cells (5 day exposure) and compared to ISO.

**Extended Data Fig. 9.**
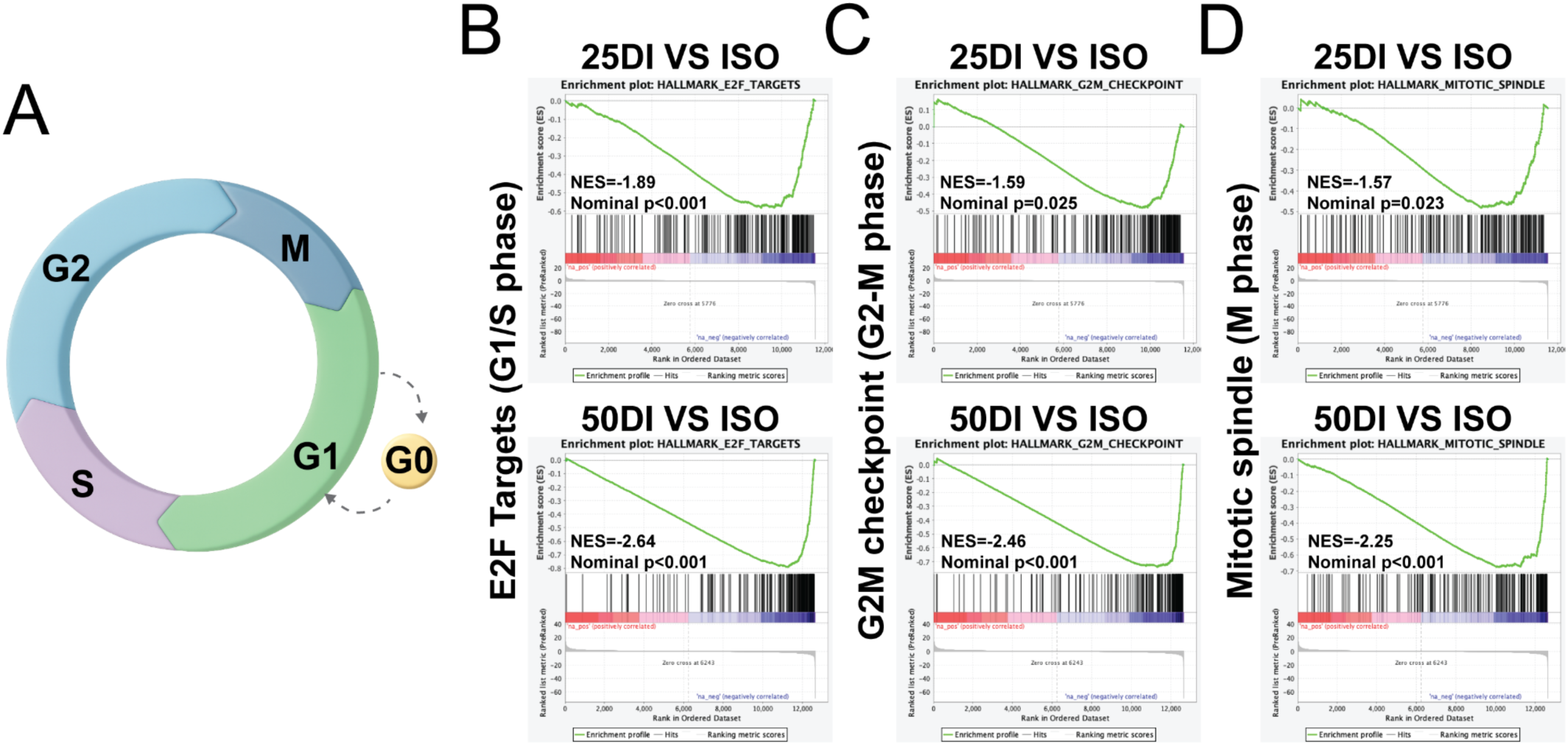
Hallmark cell cycle programs and mitotic modules confirm stronger repression in adapted 50DI. (A) Schematic of G0, G1, S, G2, and M phases of the cell cycle, shown for reference in interpreting phase grouped marker panels. (B–D) GSEA plots for Hallmark E2F targets (B), G2M checkpoint (C), and mitotic spindle (D) in adapted 25DI (upper) and adapted 50DI (lower) relative to ISO. All RNA sequencing analyses were performed on adapted cells (5 day exposure) and compared to ISO.

**Extended Data Fig. 10.**
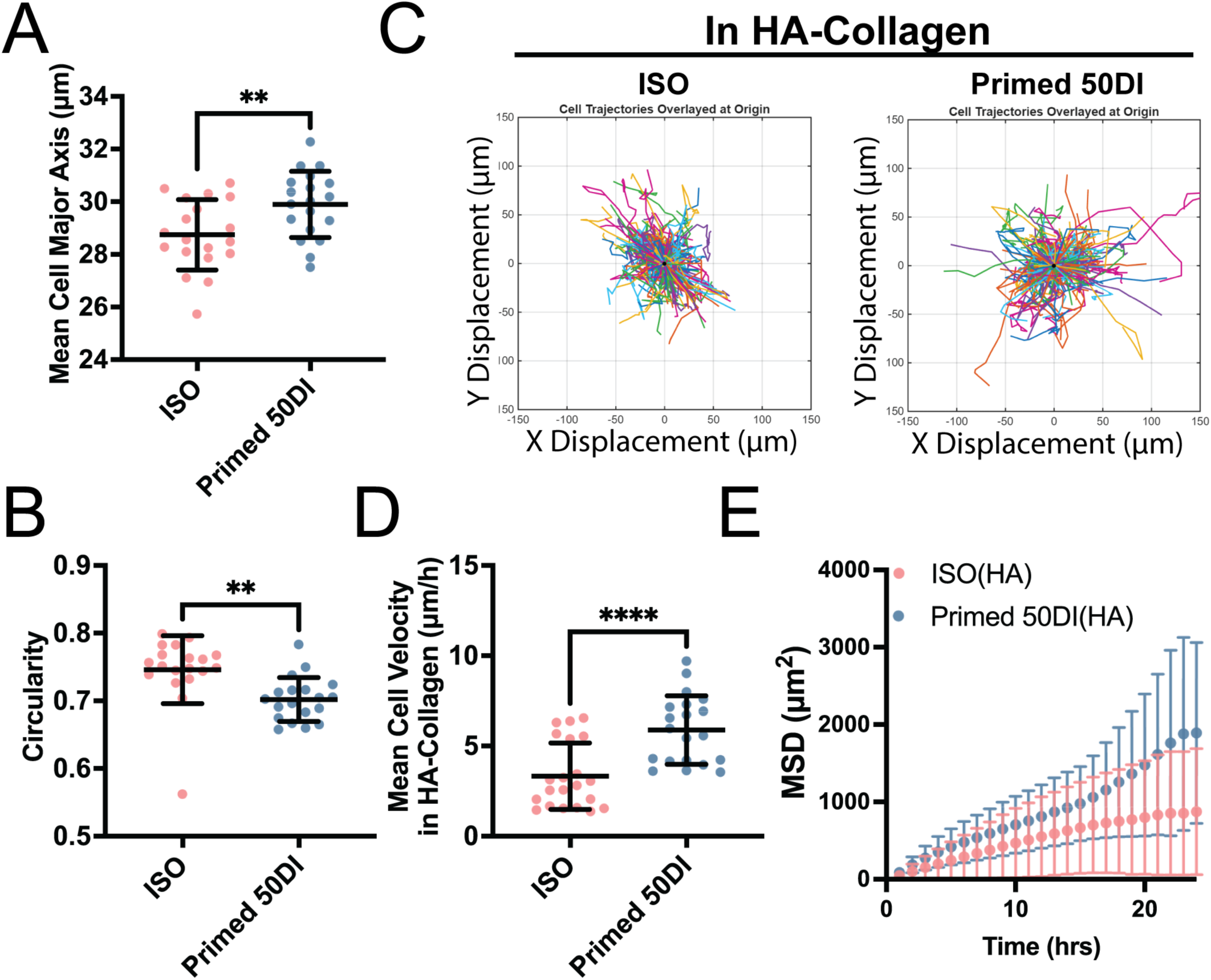
Enhanced displacement dynamics after volumetric priming are preserved in hyaluronan containing collagen matrices. (A–B) Single cell morphology in HA collagen showing mean cell major axis length (A) and circularity (B) for ISO and primed 50DI. (C) Representative x–y trajectories overlaid at the origin for ISO and primed 50DI in HA collagen. (D) Single cell mean migration velocity in HA collagen for ISO and primed 50DI. (E) Mean squared displacement over time in HA collagen for ISO and primed 50DI (n≥19 cells per condition from 4 independent experiments). Scatter plots and curves show mean ± s.d.

**Extended Data Fig. 11.**
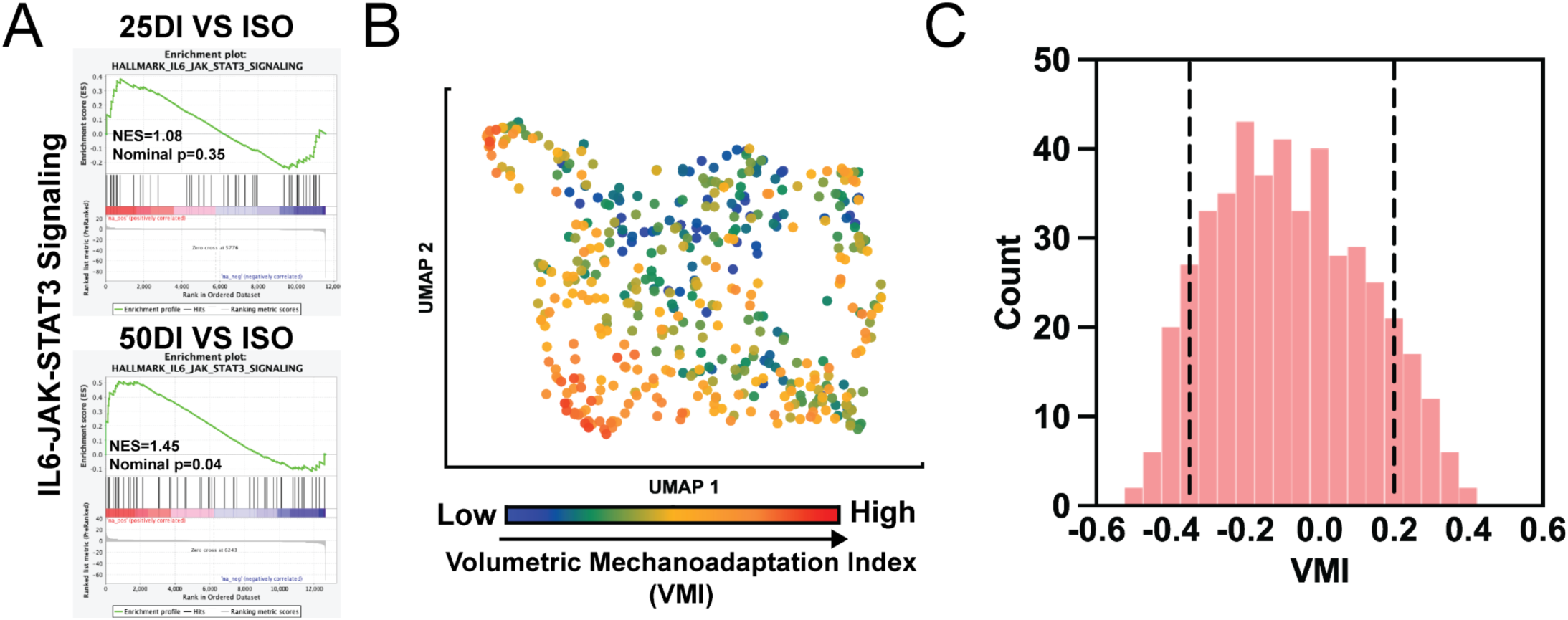
Multi-scale validation of resistance programs and construction of a patient-level VMI axis. (A) GSEA for the Hallmark IL6-JAK-STAT3 signaling gene set in adapted 25DI (upper) and adapted 50DI (lower) relative to ISO. (B) UMAP of TCGA SKCM tumors colored by Volumetric Mechanoadaptation Index (VMI), which scores similarity to the adapted 50DI transcriptional state (n=457). (C) VMI distribution across TCGA SKCM tumors, with dashed lines indicating the top 10% and bottom 10% cutoffs used for Kaplan-Meier stratification. Scatter plots show mean ± s.d. All RNA sequencing analyses were performed on adapted cells (5 day exposure) and compared to ISO.

**Extended Data Fig. 12.**
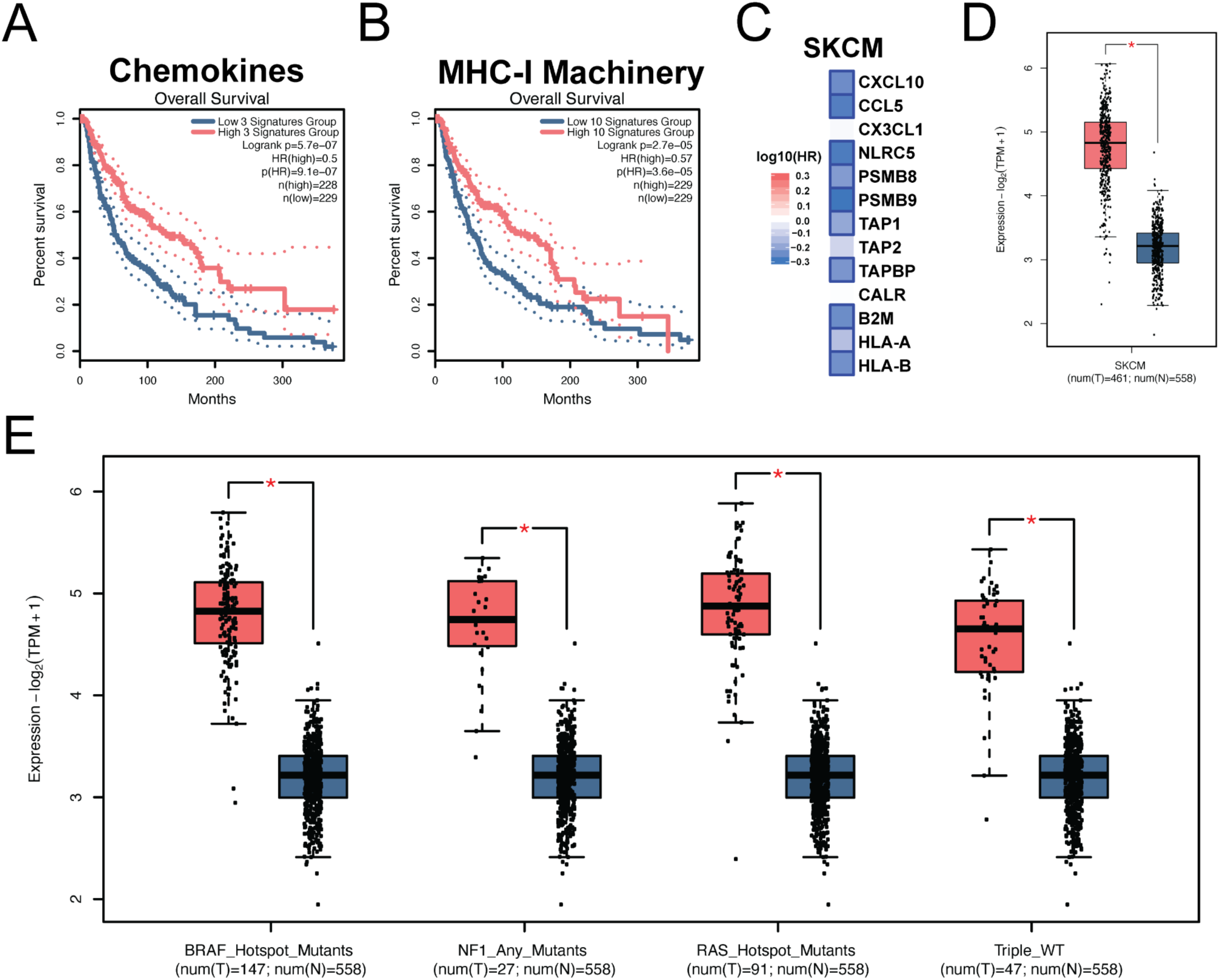
Clinical analyses support an immune engaged transcriptional axis and tumor associated sialylation programs. (A) Kaplan-Meier overall survival curves in TCGA SKCM comparing tumors with high and low expression of a chemokine module defined by CXCL10, CCL5, and CX3CL1. High and low groups were defined by the top and bottom deciles of the module score (n1=228, n2=229). (B) Kaplan-Meier overall survival curves in TCGA SKCM comparing tumors with high and low expression of an MHC-I antigen processing and presentation machinery module defined by the genes shown in Fig. 7B, using the same top and bottom decile stratification (n1=229, n2=229). (C) Summary heatmap of gene level survival associations for the chemokine and MHC-I machinery genes in Fig. 7C across TCGA SKCM. Colors indicate log10(HR) direction and magnitude, with significance indicated as squared outline of the corresponding heatmap. (D) Box plot comparing the sialic acid supply and transferase module score between TCGA SKCM tumors and non-tumor controls (n as indicated in the panel). (E) Box plot comparing the sialic acid supply and transferase module score across TCGA SKCM genomic subtypes, including BRAF_Hotspot_Mutants (n=149), RAS_Hotspot_Mutants (n=92), NF1_Any_Mutants (n=27), and Triple_WT (n=45). Non-tumor controls are shown for reference where available. All RNA sequencing analyses were performed on adapted cells (5 day exposure) and compared to ISO.

## References

1. Rambow, F. et al. Toward Minimal Residual Disease-Directed Therapy in Melanoma. Cell 174, 843–855.e19 (2018).

2. Verfaillie, A. et al. Decoding the regulatory landscape of melanoma reveals TEADS as regulators of the invasive cell state. Nat Commun 6, 6683 (2015).

3. Hunter, M. V. et al. Mechanical confinement governs phenotypic plasticity in melanoma. Nature 647, 517–527 (2025).

4. Huang, F., Santinon, F., Flores González, R. E. & Del Rincón, S. V. Melanoma Plasticity: Promoter of Metastasis and Resistance to Therapy. Front Oncol 11, 756001 (2021).

5. Puzanov, I. et al. Association of BRAF V600E/K Mutation Status and Prior BRAF/MEK Inhibition With Pembrolizumab Outcomes in Advanced Melanoma: Pooled Analysis of 3 Clinical Trials. JAMA Oncol 6, 1256–1264 (2020).

6. Grätz, V., Zillikens, D., Busch, H., Langan, E. A. & Terheyden, P. Sequential Treatment With Targeted and Immune Checkpoint Therapy in Patients With BRAF Positive Metastatic Melanoma: The Importance of Timing? Front Med (Lausanne) 6, 257 (2019).

7. Seitter, S. J. et al. Impact of Prior Treatment on the Efficacy of Adoptive Transfer of Tumor-Infiltrating Lymphocytes in Patients with Metastatic Melanoma. Clin Cancer Res 27, 5289–5298 (2021).

8. Rohaan, M. W. et al. Tumor-Infiltrating Lymphocyte Therapy or Ipilimumab in Advanced Melanoma. N Engl J Med 387, 2113–2125 (2022).

9. Nia, H. T., Munn, L. L. & Jain, R. K. Physical traits of cancer. Science 370, (2020).

10. Nia, H. T., Munn, L. L. & Jain, R. K. Probing the physical hallmarks of cancer. Nat Methods 22, 1800–1818 (2025).

11. Nia, H. T., et al. Solid stress and elastic energy as measures of tumor mechanopathology. Nat Biomed Eng 1, (2016).

12. Linke, J. A., Munn, L. L. & Jain, R. K. Compressive stresses in cancer: characterization and implications for tumor progression and treatment. Nat Rev Cancer 24, 768–791 (2024).

13. Nia, H. T. et al. Solid Stress Estimations via Intraoperative 3D Navigation in Patients with Brain Tumors. Clin Cancer Res 31, 3571–3580 (2025).

14. Stroka, K. M. et al. Water permeation drives tumor cell migration in confined microenvironments. Cell 157, 611–623 (2014).

15. Han, Y. L. et al. Cell swelling, softening and invasion in a three-dimensional breast cancer model. Nat Phys 16, 101–108 (2020).

16. McEvoy, E., Han, Y. L., Guo, M. & Shenoy, V. B. Gap junctions amplify spatial variations in cell volume in proliferating tumor spheroids. Nat. Commun. 11, 6148 (2020).

17. Chang, J. et al. Cell volume expansion and local contractility drive collective invasion of the basement membrane in breast cancer. Nat Mater 23, 711–722 (2024).

18. Guo, M. et al. Cell volume change through water efflux impacts cell stiffness and stem cell fate. Proc. Natl. Acad. Sci. U. S. A. 114, E8618–E8627 (2017).

19. Li, Y. et al. Volumetric Compression Induces Intracellular Crowding to Control Intestinal Organoid Growth via Wnt/β-Catenin Signaling. Cell Stem Cell 28, 170–172 (2021).

20. Lee, J., Abdeen, A. A., Wycislo, K. L., Fan, T. M. & Kilian, K. A. Interfacial geometry dictates cancer cell tumorigenicity. Nat Mater 15, 856–862 (2016).

21. Zhang, X. et al. Compression drives diverse transcriptomic and phenotypic adaptations in melanoma. Proc Natl Acad Sci U S A 120, e2220062120 (2023).

22. Zhao, X., Hu, J., Li, Y. & Guo, M. Volumetric compression develops noise-driven single-cell heterogeneity. Proc Natl Acad Sci U S A 118, (2021).

23. Castro-Pérez, E., Singh, M., Sadangi, S., Mela-Sánchez, C. & Setaluri, V. Connecting the dots: Melanoma cell of origin, tumor cell plasticity, trans-differentiation, and drug resistance. Pigment Cell Melanoma Res 36, 330–347 (2023).

24. Jevtić, P., Edens, L. J., Vuković, L. D. & Levy, D. L. Sizing and shaping the nucleus: mechanisms and significance. Curr Opin Cell Biol 28, 16–27 (2014).

25. Pachitariu, M., Rariden, M. & Stringer, C. Cellpose-SAM: superhuman generalization for cellular segmentation. bioRxiv (2025) doi:10.1101/2025.04.28.651001.

26. Stringer, C. & Pachitariu, M. Cellpose3: one-click image restoration for improved cellular segmentation. Nat Methods 22, 592–599 (2025).

27. Venkatareddy, M. et al. Estimating podocyte number and density using a single histologic section. J Am Soc Nephrol 25, 1118–1129 (2014).

28. Gut, G., Tadmor, M. D., Pe’er, D., Pelkmans, L. & Liberali, P. Trajectories of cell-cycle progression from fixed cell populations. Nat Methods 12, 951–954 (2015).

29. Zhang, S. et al. Generation of cancer stem-like cells through the formation of polyploid giant cancer cells. Oncogene 33, 116–128 (2014).

30. Moran, P. A. P. Notes on continuous stochastic phenomena. Biometrika 37, 17–23 (1950).

31. Anselin, L. Local indicators of spatial association—LISA. Geogr. Anal. 27, 93–115 (1995).

32. Lee, H.-P., Stowers, R. & Chaudhuri, O. Volume expansion and TRPV4 activation regulate stem cell fate in three-dimensional microenvironments. Nat. Commun. 10, 529 (2019).

33. Hänzelmann, S., Castelo, R. & Guinney, J. GSVA: gene set variation analysis for microarray and RNA-seq data. BMC Bioinformatics 14, 7 (2013).

34. Liberzon, A. et al. The Molecular Signatures Database (MSigDB) hallmark gene set collection. Cell Syst 1, 417–425 (2015).

35. Subramanian, A. et al. Gene set enrichment analysis: a knowledge-based approach for interpreting genome-wide expression profiles. Proc Natl Acad Sci U S A 102, 15545–15550 (2005).

36. Denais, C. M. et al. Nuclear envelope rupture and repair during cancer cell migration. Science 352, 353–358 (2016).

37. Irianto, J., Lee, D. A. & Knight, M. M. Quantification of chromatin condensation level by image processing. Med Eng Phys 36, 412–417 (2014).

38. Na, J. et al. Mechanical memory based on chromatin and metabolism remodeling promotes proliferation and smooth muscle differentiation in mesenchymal stem cells. FASEB J 38, e23538 (2024).

39. Shi, T. & Dansen, T. B. Reactive Oxygen Species Induced p53 Activation: DNA Damage, Redox Signaling, or Both? Antioxid Redox Signal 33, 839–859 (2020).

40. Ji, S. et al. Baf60b-mediated ATM-p53 activation blocks cell identity conversion by sensing chromatin opening. Cell Res 27, 642–656 (2017).

41. Hoek, K. S. et al. In vivo switching of human melanoma cells between proliferative and invasive states. Cancer Res 68, 650–656 (2008).

42. Eichhoff, O. M. et al. The immunohistochemistry of invasive and proliferative phenotype switching in melanoma: a case report. Melanoma Res 20, 349–355 (2010).

43. Schwarzbauer, J. E. & DeSimone, D. W. Fibronectins, their fibrillogenesis, and in vivo functions. Cold Spring Harb Perspect Biol 3, (2011).

44. Lu, P., Weaver, V. M. & Werb, Z. The extracellular matrix: a dynamic niche in cancer progression. J Cell Biol 196, 395–406 (2012).

45. Dalgleish, R. The human type I collagen mutation database. Nucleic Acids Res 25, 181–187 (1997).

46. Te Molder, L., de Pereda, J. M. & Sonnenberg, A. Regulation of hemidesmosome dynamics and cell signaling by integrin α6β4. J Cell Sci 134, (2021).

47. Sil, H., Sen, T. & Chatterjee, A. Fibronectin-integrin (alpha5beta1) modulates migration and invasion of murine melanoma cell line B16F10 by involving MMP-9. Oncol Res 19, 335–348 (2011).

48. McKenzie, J. A., Liu, T., Goodson, A. G. & Grossman, D. Survivin enhances motility of melanoma cells by supporting Akt activation and {alpha}5 integrin upregulation. Cancer Res 70, 7927–7937 (2010).

49. Etienne-Manneville, S. & Hall, A. Rho GTPases in cell biology. Nature 420, 629–635 (2002).

50. Lee, H. W. et al. Alpha-smooth muscle actin (ACTA2) is required for metastatic potential of human lung adenocarcinoma. Clin Cancer Res 19, 5879–5889 (2013).

51. Gao, S. et al. TUBB4A interacts with MYH9 to protect the nucleus during cell migration and promotes prostate cancer via GSK3β/β-catenin signalling. Nat Commun 13, 2792 (2022).

52. Onder, T. T. et al. Loss of E-cadherin promotes metastasis via multiple downstream transcriptional pathways. Cancer Res 68, 3645–3654 (2008).

53. Zhang, X. & Mak, M. Biophysical Informatics Approach For Quantifying Phenotypic Heterogeneity In Cancer Cell Migration In Confined Microenvironments. Bioinformatics 37, 2042–2052 (2021).

54. Wisdom, K. M. et al. Matrix mechanical plasticity regulates cancer cell migration through confining microenvironments. Nat Commun 9, 4144 (2018).

55. Han, Y. et al. Long noncoding RNA LINC00239 inhibits ferroptosis in colorectal cancer by binding to Keap1 to stabilize Nrf2. Cell Death Dis 13, 742 (2022).

56. Sherwood, A. M., Yasseen, B. A., DeBlasi, J. M., Caldwell, S. & DeNicola, G. M. Distinct roles for the thioredoxin and glutathione antioxidant systems in Nrf2-Mediated lung tumor initiation and progression. Redox Biol 83, 103653 (2025).

57. Shi, X. et al. Huaier suppresses lung cancer by simultaneously and independently inhibiting the antioxidant pathway SLC7A11/GPX4 while enhancing ferritinophagy. Cell Death Discov 11, 309 (2025).

58. Ara, T. et al. Critical role of STAT3 in IL-6-mediated drug resistance in human neuroblastoma. Cancer Res 73, 3852–3864 (2013).

59. Tian, X. et al. Targeting apoptotic pathways for cancer therapy. J Clin Invest 134, (2024).

60. Stifter, K. et al. IFN-γ treatment protocol for MHC-I/PD-L1 pancreatic tumor cells selectively restores their TAP-mediated presentation competence and CD8 T-cell priming potential. J Immunother Cancer 8, (2020).

61. Gray, M. A. et al. Targeted glycan degradation potentiates the anticancer immune response in vivo. Nat Chem Biol 16, 1376–1384 (2020).

62. Wu, J. et al. Targeted glycan degradation potentiates cellular immunotherapy for solid tumors. Proc Natl Acad Sci U S A 120, e2300366120 (2023).

63. Park, S. et al. Immunoengineering can overcome the glycocalyx armour of cancer cells. Nat Mater 23, 429–438 (2024).

64. Yang, Z. et al. Targeted desialylation and cytolysis of tumor cells by fusing a sialidase to a bispecific T-cell engager. Nat Biomed Eng 8, 499–512 (2024).

65. Foote, C. A. et al. Neuraminidase inhibition improves endothelial function in diabetic mice. Am J Physiol Heart Circ Physiol 325, H1337–H1353 (2023).

66. Bera, K. et al. Extracellular fluid viscosity enhances cell migration and cancer dissemination. Nature 611, 365–373 (2022).

67. Engeland, K. Cell cycle arrest through indirect transcriptional repression by p53: I have a DREAM. Cell Death Differ 25, 114–132 (2018).

68. Palmer, A. C., Chidley, C. & Sorger, P. K. A curative combination cancer therapy achieves high fractional cell killing through low cross-resistance and drug additivity. Elife 8, (2019).

69. Puig, I. et al. TET2 controls chemoresistant slow-cycling cancer cell survival and tumor recurrence. J Clin Invest 128, 3887–3905 (2018).

70. Carlson, P. et al. Targeting the perivascular niche sensitizes disseminated tumor cells to chemotherapy. Nat Cell Biol 21, 238–250 (2019).

71. Robert, C. et al. Ipilimumab plus dacarbazine for previously untreated metastatic melanoma. N Engl J Med 364, 2517–2526 (2011).

72. Dutta, S. & Sen, S. Preparation and Characterization of Collagen-Hyaluronic Acid (Col-HA) Matrices: In Vitro Mimics of the Tumor Microenvironment. Methods Mol Biol 2747, 131–139 (2024).

73. Tang, Z., Kang, B., Li, C., Chen, T. & Zhang, Z. GEPIA2: an enhanced web server for large-scale expression profiling and interactive analysis. Nucleic Acids Res 47, W556–W560 (2019).

74. Shi, J. et al. The Web-Based Portal SpatialTME Integrates Histological Images with Single-Cell and Spatial Transcriptomics to Explore the Tumor Microenvironment. Cancer Res 84, 1210–1220 (2024).

